# Genetic identification of the functional surface for RNA binding by *Escherichia coli* ProQ

**DOI:** 10.1101/822791

**Authors:** Smriti Pandey, Chandra M. Gravel, Oliver M. Stockert, Clara D. Wang, Courtney L. Hegner, Hannah LeBlanc, Katherine E. Berry

**Author notes:** To whom correspondence should be addressed. KB. [Smriti Pandey], Biological and Biomedical Sciences Program, Harvard University, Boston, MA, 02115, USA. [Clara D. Wang]: National Heart, Lung, and Blood Institute, National Institutes of Health, Bethesda, MD, 20892, USA. [Courtney L. Hegner]: Skaggs Graduate Program, Scripps Research Institute, Jupiter, FL, 33458, USA.

## Abstract

The FinO-domain-protein ProQ is an RNA-binding protein that has been known to play a role in osmoregulation in proteobacteria. Recently, ProQ has been shown to act as a global RNA-binding protein in *Salmonella* and *E. coli*, binding to dozens of small RNAs (sRNAs) and messenger RNAs (mRNAs) to regulate mRNA-expression levels through interactions with both 5’ and 3’ untranslated regions (UTRs). Despite excitement around ProQ as a novel global RNA-binding protein interacting with many sRNAs and mRNAs, and its potential to serve as a matchmaking RNA chaperone, significant gaps remain in our understanding of the molecular mechanisms ProQ uses to interact with RNA. In order to apply the tools of molecular genetics to this question, we have adapted a bacterial three-hybrid (B3H) assay to detect ProQ’s interactions with target RNAs. Using domain truncations, site-directed mutagenesis and an unbiased forward genetic screen, we have identified a group of highly conserved residues on ProQ’s NTD as the primary face for *in vivo* recognition of two RNAs, and propose that the NTD structure serves as an electrostatic scaffold to recognize the shape of an A-form RNA duplex.

## INTRODUCTION

Regulatory, small RNAs (sRNAs) are found in nearly all bacterial species and implicated in important processes such as virulence, biofilm formation, host interactions and antibiotic resistance.(1-3) These sRNAs typically regulate messenger RNA (mRNA) translation through imperfect base pairing near an mRNA’s ribosomal binding site.(2, 4-6) In many bacterial species, the stability and function of sRNAs are supported by global RNA-binding proteins, such as the protein Hfq.(1, 4, 7-9) Given that Hfq is not present in all bacterial species and that not all sRNAs depend on Hfq for their function, there is increasing interest in other RNA-binding proteins that may play a role in global gene-regulation in bacteria,(2, 10-13) including a class of proteins that contain FinO domains.(14-17)

The *Escherichia coli* protein FinO is the founding member of the FinO structural class of RNA-binding proteins. In *E. coli*, FinO binds the FinP sRNA and regulates the 5’ untranslated region (UTR) of *traJ*.(18, 19) Similarly, *Legionella pneumophila* RocC contains a FinO-domain and binds the sRNA RocR along with at least four 5’ UTRs of mRNAs involved in competence.(20) In *E. coli*, another FinO-domain-containing protein called ProQ was initially characterized as an RNA-binding protein contributing to osmoregulation through expression of *proP*.(21) ProQ was recently identified through Grad-Seq experiments to bind to dozens of cellular RNAs,(17) including a large number of sRNAs and mRNA 3’UTRs in *Samonella* and *E. coli*.(22) ProQ binding has been shown to regulate mRNA-expression levels through interactions with both 5’ and 3’ UTRs. It has been shown to form a ternary complex with an sRNA (RaiZ) and an mRNA (*hupA*), to support RaiZ’s repression of *hupA*,(23) and to protect mRNAs from exonucleolytic degradation by binding to 3’ ends.(22) Further, ProQ supports the sRNA SraL in preventing premature termination of *rho* transcripts in *Salmonella*,(24) and promotes *Salmonella* invasion of HeLa cells.(25) Global analysis of ProQ-bound RNAs using UV CLIP-seq suggests that ProQ interacts with highly structured RNAs, with a simple 12-bp hairpin as the consensus motif.(22) This is consistent with *in vitro* analysis showing that FinO’s binding affinity for FinP RNA depends on the presence of an RNA duplex rather than the sequence of bases within the duplex, and that FinO protects the base of RNA duplex stems and the nucleotides immediately 3’ of the stem.(26, 27) However, the specific determinants of ProQ’s binding preferences for cellular RNAs have yet to be determined.

ProQ’s domain architecture consists of structured N-terminal and C-terminal domains (NTD, CTD) with a poorly conserved and likely unstructured linker connecting them (Fig S1). NMR structures for both conserved domains of *E. coli* ProQ have been solved, demonstrating that the NTD adopts a FinO-like fold.(28) RNA-binding studies have offered conflicting information about the domain(s) responsible for RNA binding: the NTD/FinO-domain of ProQ has been shown to be sufficient for high-affinity binding to dsRNA *in vitro*,(21) as has the FinO-domain of RocC for high affinity binding to the RocR sRNA.(20) On the other hand, biophysical data indicate that the chemical environment of residues both in the NTD and also in the linker and CTD change in the presence of RNA substrates.(28) Within the NTD/FinO-domain of ProQ, a crosslinking study found that lysine and arginine residues on both surfaces of structure contacted RNA,(29) while biophysical experiments implicate one face more than the other RNA binding.(28) Thus, there is still significant uncertainty about the functional RNA-binding domains and surfaces of ProQ. Critically, there has been no comprehensive mutagenesis conducted to map the functional binding surface of ProQ in recognizing its sRNA and mRNA substrates.

We have previously reported a transcription-based bacterial three-hybrid (B3H) assay that facilitates the detection of RNA-protein interactions inside of living *E. coli* reporter cells.(30) While this assay was effective in detecting numerous Hfq-sRNA interactions, it was unclear how generally applicable this assay would be to other RNA-protein interactions. Here we present a modified B3H assay that is able to robustly and specifically detect ProQ-RNA interactions, and provides a path to apply the tools of molecular genetics to the mechanism of ProQ-RNA interactions. We utilize this assay as a platform for targeted mutation of highly conserved residues as well as an unbiased forward genetic screen to define the functional RNA-binding surface of ProQ. We have identified multiple single-point mutations that disrupt ProQ’s interaction with target RNAs. Our data suggest that the conserved N-terminal FinO-domain is the principal site of RNA binding *in vivo* for both an sRNA and 3’UTR. Using available NMR structures for the ProQ NTD, and guided by the results of our forward and reverse genetic approaches, we present a working model for molecular recognition between ProQ and interacting RNAs. We demonstrate the necessity of more than eight residues across a highly conserved face of the NTD for strong RNA binding by ProQ. The chemical nature and location of these residues suggest that ProQ uses a combination of electrostatic, hydrogen-bonding and hydrophobic interactions over a large surface area to mediate RNA interactions. We propose that ProQ achieves specificity for duplex RNA by acting as an electrostatic scaffold, with the overall structure of the NTD/FinO-domain serving to position several charged residues in an appropriate geometry to read out the shape of an A-form RNA double helix.

## MATERIAL AND METHODS

### Bacterial strains and plasmids

A complete list of plasmids, strains, and oligonucleotides (oligos) used in this study is provided in Supplemental Tables S1-S3, respectively. NEB 5-alpha F’Iq cells (New England Biolabs) were used as the recipient strain for all plasmid constructions.

A single-copy O_L_2-62-*lacZ* reporter on an F’ episome bearing tetracycline resistance was generated as previously described (31, 32) by conjugative delivery of pFW11-derivative plasmid pFW11-O_L_2-tet into FW102 cells to generate *Escherichia coli* strain KB480, which is analogous to O_L_2-62-*lacZ* reporters carried on F’ episomes bearing kanamycin resistance.(30, 33) The *Δhfq::kan* allele and *ΔproQ::kan* allele from the Keio collection(34) were introduced to KB480 via P1 transduction to generate KB483 and SP2 respectively. An analogous process was used to create strain KB511 from by conjugative delivery of pFW11-derivative plasmid pKB1067 into FW102 cells. pKB1067 was generated from pFW11_tet_ O_L_2-62-*lacZ* from overlap PCR with oKB1366 + oKB1367 to insert a 21bp-sequence (GCTGCCACGGTGCCCGACCGT) immediately downstream of O_L_2 site. Thus, KB511 carries a single copy *O*_*L*_*2-83-lacZ* reporter on F’ episome bearing tetracycline resistance, in which the lambda operator *O*_*L*_*2* is centered at a position of −83 relative to the transcription start site (TSS). The recombinant F’ episome was then moved via conjugation into *Δhfq* strain KB496 to give strain SP5, which was used as the reporter strain for the unbiased screen (described below). Except for this screen and data presented in Figure 1, KB483 (*O*_*L*_*2-62-lacZ; Δhfq*) was used as the reporter strain for all B3H experiments in this study.

**Figure 1.**
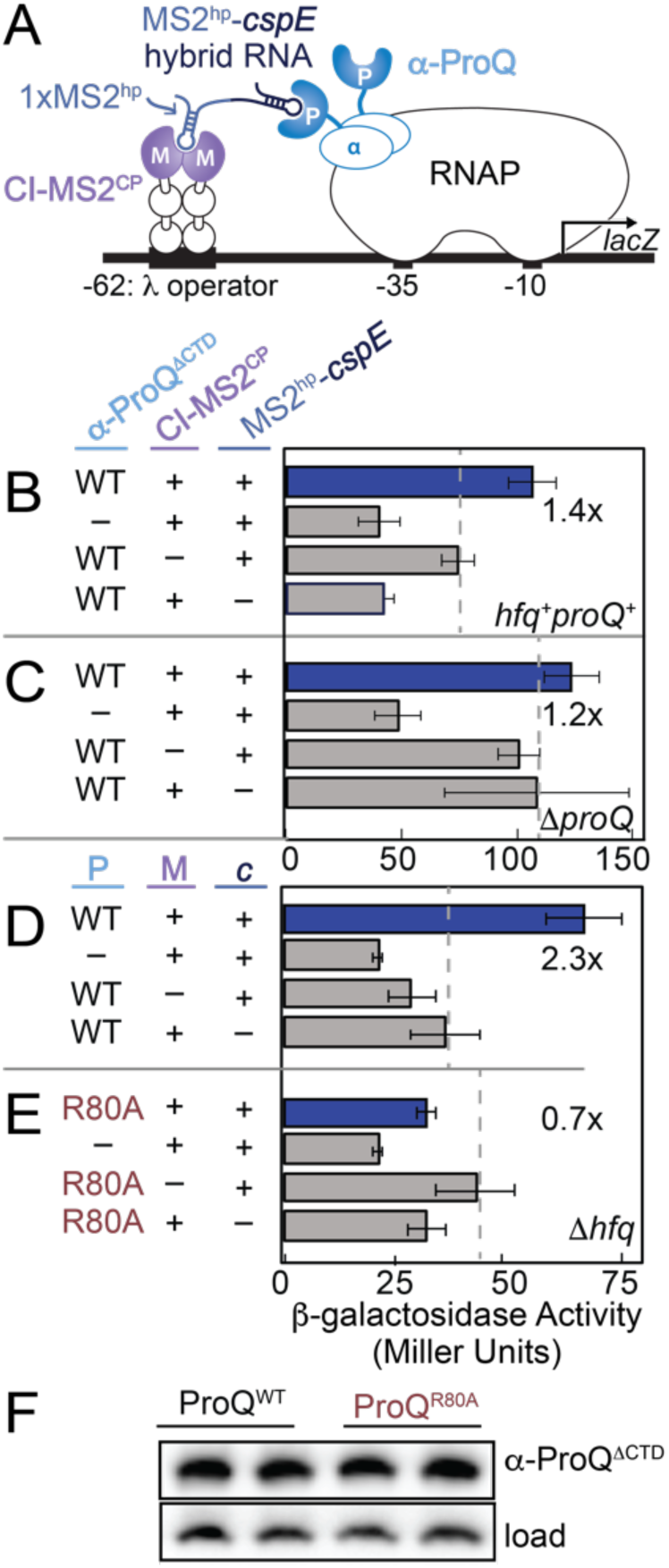
Adaptation of an *E. coli* bacterial three-hybrid (B3H) interaction to detect ProQ-RNA interactions. (A) Design of B3H system to detect interaction between ProQ and an RNA (*cspE* 3’UTR). Interaction between protein moiety ProQ and RNA moiety *cspE* fused, respectively, to the α subunit of RNAP (α-NTD) and to one copy of the MS2 RNA hairpin (MS2^hp^) activates transcription from test promoter, which directs transcription of a *lacZ* reporter gene. The test promoter (p*lac*-O_L_2–62), which bears the λ operator O_L_2 centered at position –62 relative to the transcription start site, is present on a single copy F’ episome (33). The RNA-binding moiety MS2^CP^ is fused to λCI (CI-MS2^CP^) to tether the hybrid RNA (MS2^hp^-*cspE*) to the test promoter. Compatible plasmids direct the synthesis of the α-fusion protein (under the control of an IPTG-inducible promoter), the CI-MS2^CP^ adapter protein (under the control of a constitutive promoter; pCW17) and the hybrid RNA (under the control of an arabinose-inducible promoter). (B-E) Results of β-galactosidase assays performed with wild type (B; *hfq*^*+*^*proQ*^*+*^), Δ*proQ* (C) or Δ*hfq* (D,E) reporter strain cells containing three compatible plasmids: one (α-ProQ; P) that encoded α (–) or the α-ProQ^ΔCTD^ (pKB955; resi=2-176) fusion protein (WT or an R80A mutant), another (CI-MS2^CP^; M) that encoded λCI (–) or the λCI-MS2^CP^ fusion protein (+), and a third (MS2^hp^-*cspE, c*) that encoded a hybrid RNA with the 3’ UTR of *cspE* (pSP10, final 85 nts) following one copy of an MS2^hp^ moiety (+) or an RNA that contained only the MS2^hp^ moiety (–). Cells were grown in the presence of 0.2% arabinose and 50 μM IPTG (see Methods). All subsequent assays were performed in Δ*hfq* reporter strain cells. Bar graphs show the averages of three independent measurements and standard deviations. (F) Samples from (D) and (E) were analyzed by Western blot and probed with an anti-ProQ antibody detect α-ProQ^ΔCTD^ fusion protein (α-ProQ). A cross-reacting band independent of the presence of endogenous ProQ or α-ProQ fusion protein is used as a loading control (load; see Fig S2). Duplicate biological samples are shown.

Plasmids were constructed as specified in Table S1. PCR mutagenesis to create site-directed mutants of *proQ* was conducted with the Q5 Site-Directed Mutagenesis Kit (New England Biolabs) using end-to-end primers designed with NEBaseChanger. The construction of key parent vectors is described below. Residues 2-119, 2-131, 2-176, 181-232 and 2-232 of *E. coli* ProQ were fused to the α-NTD (residues 1-248) in pBRα between NotI and BamHI to generate pSP90 (pBRα-ProQ^NTD^), pKB951 (pBRα-ProQ^NTD+12aa^), pKB955 (pBRα-ProQ^ΔCTD^), pSP92 (pBRα-ProQ^CTD^) and pKB949 (pBRα-ProQ^FL^, full-length) respectively.

pCW17 (pAC-p_constit_-λCI-MS2^CP^) was derived from pKB989 (pAC-p_lacUV5_-λCI-MS2^CP^) (22) by substitution of the region between −35 and +22 of the p_lacUV5_ promoter (containing the −35, −10 and *lacO* elements) with the following sequence lacking a *lacO* element (predicted −35, −10 and TSS of the resulting constitutive promoter are underlined):

CTCGAG<u>ACGATA</u>GCTAGCTCAGTCCTAGG<u>TATAGT</u>GCTAGC<u>G</u>CATGC Stepwise, this substitution was made using Gibson Assembly of a backbone (PCR product of oCW6 and oCW7 on pKB989) and insert (hybridization of oCW18 and oCW19), followed by mutagenesis PCR on the resulting plasmid (with primers oCW28 and oCW29).

pCH1 (pCDF-pBAD-1xMS2^hp^-XmaI-HindIII) was derived from pKB845 (pCDF-pBAD-2x MS2^hp^-XmaI-HindIII),(30) by removing 2xMS2^hp^ moieties via vector digestion (EcoRI + XmaI) followed by ligation of an insert formed by kinase-treated oCH1 + oCH2, which encodes an EcoRI site, one copy of a 21-nt RNA hairpin from bacteriophage MS2 (MS2^hp^), and an XmaI site. All hybrid RNAs used in this study are 1xMS2^hp^-RNA hybrids and constructed by inserting the RNA of interest into the XmaI/HindIII sites of pCH1.

### β-galactosidase assays

For B3H assays, reporter cells (KB480, KB483 or SP2) were freshly co-transformed with compatible pAC-, pBR- and pCDF-derived plasmids, as indicated. From each transformation three colonies (unless otherwise noted) were picked into 1 ml LB broth supplemented with carbenicillin (100 μg/ml), chloramphenicol (25 μg/ml), tetracycline (10 μg/ml), spectinomycin (100 μg/ml) and 0.2% arabinose in a 2 ml 96-well deep well block (VWR), sealed with breathable film (VWR) and shaken at 900 rpm at 37°C. Overnight cultures were diluted 1:50 into 200 μl LB supplemented as above, with an additional 50 μM isopropyl-β-D-thiogalactoside (IPTG) when noted. Cells were grown to mid-log (OD_600_≈0.6) in optically clear 200 μl flat bottom 96-well plates (Olympus) covered with plastic lids, as above. Cells were lysed and β-galactosidase (β-gal) activity was measured as previously described.(35) B3H interactions are calculated and reported as the fold-stimulation over basal levels; this is the β-gal activity in reporter cells containing all hybrid constructs (α-ProQ, CI-MS2^CP^ and MS2^hp^-Bait-RNA), divided by the highest activity from negative controls—cells containing plasmids where one of the hybrid constructs is replaced by an α empty, CI empty or MS2^hp^ empty construct. Assays were conducted in biological triplicate on at least three separate days. Absolute β-gal values from a representative dataset of a biological triplicate experiment, including values for all negative controls, are shown in Supplementary Figures 4 and 6 as mean β-gal values arising from one triplicate experiment. In main-text figures, B3H interactions are reported as average values of fold-stimulation over basal levels from at least three experiments across multiple days, and the standard deviation of these average values from multiple independent experiments.

### Western Blots

Cell lysates from β-gal assays were normalized based on pre-lysis OD_600_. Lysates were mixed with 6× Laemmli loading dye with PopCulture Reagent (EMD Millipore Corp), boiled for 10 min at 95°C and electrophoresed on 10–20% Tris-glycine gels (Thermo Fisher) in 1x NuPAGE MES Running Buffer (Thermo Fisher). Proteins were transferred to PVDF membranes (BioRad) using a semi-dry transfer system (BioRad Trans-blot Semi-Dry and Turbo Transfer System) according to manufacturer’s instructions, and probed with 1:10,000 primary antibody (anti-RpoA-NTD; Neoclone or anti-ProQ; kindly provided by G. Storz) overnight at 4°C and then a horseradish peroxidase (HRP)-conjugated secondary antibody (anti-mouse IgG or anti-rabbit IgG; Cell Signaling, 1:10,000). Note that, throughout the paper, “anti-ProQ” is written out rather than using the standard abbreviation of “α-ProQ.” This is to avoid confusion with the fusion protein we call “α-ProQ” consisting of the NTD of RpoA (α) fused in frame to ProQ. Chemiluminescent signal from bound peroxidase complexes was detected using ECL Plus western blot detection reagents (BioRad) and a c600 imaging system (Azure) according to manufacturer’s instructions.

### Random Mutagenesis

A mutant *proQ* library was generated first by 30 rounds of PCR amplification of the *proQ* portion of the pBRα-ProQ^FL^ plasmid (pKB949) using Phusion DNA Polymerase (New England Biolabs) in 70 mM MgCl_2_, 500 mM KCl, 100 mM Tris (pH 8.0), 0.1% bovine serum albumin, 2 mM dGTP, 2 mM dATP, 10 mM dCTP, 10 mM dTTP and primers oKB1077 and oKB1078. A second mutant *proQ* library was generated under the same condition but with the addition of 0.1 mM MnCl_2_.(36) The PCR products of both libraries were digested with DpnI (New England Biolabs) to remove template plasmid, then with NotI-HF and BamHI-HF (New England Biolabs), gel purified, and ligated (T4 DNA ligase; New England Biolabs) into a pBRα vector cut with NotI-HF and BamHI-HF. Following ligation and transformation into NEB 5-alpha F’Iq cells (New England Biolabs), cells were grown as near-lawns on LB-carbenicillin plates and a miniprep was performed from resuspension of ∼23,000 colonies to yield the plasmid library.

### B3H Screening and Dot Blots

For dot-blots, cell lysates (3 uL) from β-gal assays were transferred to nitrocellulose Protran membranes (Amersham) by multichannel pipette. Membranes were allowed to dry, then probed and imaged as above. To verify that the dot-blot assay could identify destabilized α-ProQ proteins, 10 *proQ* mutants were sequenced, all of which resulted in reduced β-gal activity on plates and in liquid, 5 of which showed reduced levels by dot blot and 5 of which showed approximate wild-type-levels by dot blot. There was a 100% correspondence between the levels of ProQ indicated by dot blot and the presence or absence of a premature stop codon in the NTD (Table S4).

For the primary screen, the pBRα-proQ plasmid library was transformed into SP5 cells along with pCW17 (pACλCI-MS2^CP^) and pSP10 (pCDF-MS2^hp^-*cspE*) or pSP14 (pCDF-MS2^hp^-SibB) and plated on LB agar supplemented with inducers (0.2% arabinose and 1.5 βM IPTG), antibiotics (carbenicillin (100 βg/ml), chloramphenicol (25 βg/ml), tetracycline (10 βg/ml), and spectinomycin (100 βg/ml)) and indicators (5-bromo-4-chloro-3-indolyl-β-D-galactopyranoside (Xgal; 40 βg/mL) and phenylethyl-β-D-thiogalactopyranoside (TPEG; 75 βM; Gold Biotech)). These conditions were chosen to enable a clear distinction between blue positive-control colonies (containing the WT fusion proteins and the *cspE* hybrid RNA) and white negative-control colonies (instead containing a plasmid encoding α-empty). Reporter strain SP5, (*O*_*L*_*2-83-lacZ;*) was used only for the high-throughput screen and results of RNA-binding defects were subsequently verified in KB483 (*O*_*L*_*2-62-lacZ*). Plates were incubated overnight at 37°C, then at 4°C for an additional ∼48 hours. 536 white or pale colonies were isolated (372 against *cspE* + 164 against SibB), and liquid β-gal assays were conducted to confirm the effects of these white colonies, followed by a dot-blot counter-screen with anti-ProQ antibody to eliminate mutants with low expression levels. To analyze dot-blot results of colonies that were identified in the primary screen, densitometry was conducted using ImageJ software to quantify the intensity of individual dots. The intensity of each dot was normalized to the OD_600_ of the culture before lysis and β-gal activity of each colony was plotted against normalized ProQ intensity. Normalized intensities were compared to positive and negative controls and colonies with ProQ-expression levels in the wild-type range were selected for sequencing.

Plasmids were isolated from 86 colonies that produced low β-gal activity but wild-type levels of α-ProQ fusion protein, and the DNA encoding *proQ* in each pBRα-*proQ* plasmid was sequenced. 37 colonies were found to carry pBRα-*proQ* plasmids containing single mutations that encoded unique substitutions in α-ProQ. Mini-prepped plasmids from these colonies were re-transformed into KB483 reporter cells already carrying pCW17 (pACλCI-MS2^CP^) and pSP10 (pCDF-1xMS2^hp^-*cspE*) or pSP14 (pCDF-1xMS2^hp^-SibB). Liquid β-gal assays were conducted as above, with induction of α-ProQ from both 0 βM and 50 βM IPTG, and ProQ-expression levels were evaluated in triplicate at IPTG concentrations. The basal level β-gal activity was set by activity in reporter cells containing an α-empty plasmid rather than an pBRα-*proQ* plasmid isolated in the screen. Average fold-stimulation of β-gal activity and dot-blot intensities of each mutant were then normalized to the values of WT α-ProQ (set to 1.0) and α-empty (set to 0.0) using the expression (Value_mutant_-Value_α-empty_)/(Value_WT,0 IPTG_-Value_α-empty,0 IPTG_). Note that relative expression and fold-stimulation of each mutant at 0 μM and 50 μM IPTG can be directly compared to one another, as both sets of values were normalized to WT α-ProQ at 0 μM IPTG.

## RESULTS

### Establishing a B3H assay for ProQ-RNA interactions

Our previous studies with the bacterial three-hybrid (B3H) assay have focused on Hfq-RNA interactions.(30) We sought to determine whether this B3H assay could detect interactions of ProQ with RNA in an analogous manner to the interactions of RNA and Hfq. The *cspE* 3’UTR was chosen as an initial RNA candidate due to strong interaction with ProQ that has been observed both *in vivo* and *in vitro.*(22, 28) To simplify the possibility of multiple ProQ domains interacting with RNA, we began our analysis with a construct possessing only the ProQ NTD and linker (resi 2-176; hereafter ProQ^ΔCTD^). For the envisioned assay, ProQ^ΔCTD^, the “prey” protein, is fused to the N-terminal domain of the alpha subunit of RNAP (α; Fig 1A). A single-copy test promoter contains the operator O_L_2 centered 62-bp upstream from the transcription-start site (TSS) of a *lacZ* reporter gene. The 3’-terminal 85 nts of the *cspE* transcript (hereafter *cspE*) serves as the “bait” and is expressed as a hybrid RNA following one copy of a 21-nt RNA hairpin from bacteriophage MS2 (MS2^hp^). This hybrid RNA is tethered to the upstream O_L_2 DNA sequence via a constitutively expressed RNA-DNA “adapter” protein consisting of a fusion between the CI protein from bacteriophage λ (CI) and the coat protein from bacteriophage MS2 (MS2^CP^). In this system, interaction between DNA-tethered *cspE* RNA and the RNAP-assembled α-NTD-ProQ fusion protein stabilizes the binding of RNAP to the test promoter, thereby activating reporter gene expression.

We asked whether the interaction of *cspE* with α-ProQ^ΔCTD^ would stimulate *lacZ* expression from the −62-O_L_2 test promoter (Fig 1A). β-galactosidase (β-gal) assays show that transcription from the test promoter is stimulated slightly (∼1.4-fold) when all three hybrid components are present, as compared to the basal activity from the negative controls where any single element (ProQ, *cspE*, or MS2^CP^) is left out (Fig 1B). We wondered whether the ProQ-*cspE* B3H interaction might be limited by competition with endogenous ProQ or cellular Hfq, given that co-immunoprecipitation studies have suggested that ProQ and Hfq may compete for a subset of their RNA substrates.(22, 37) While no additional stimulation of transcription over basal levels was observed when the ProQ-*cspE* B3H experiment was repeated in Δ*proQ* reporter cells (Fig 1C; 1.2), we found that the fold-stimulation of β-gal transcription indeed increased in Δ*hfq* reporter cells (Fig 1D; 2.3x). This Δ*hfq* reporter strain was previously used for detecting Hfq-RNA interactions,(30) and was used throughout the remainder of this study. To confirm the ProQ-*cspE* B3H interaction represented a specific interaction, we sought to disrupt it with a point substitution. We chose Arg80 as a conserved residue, which has previously been proposed to mediate RNA interactions,(28) to replace with alanine within the ProQ fusion protein. Unlike with WT ProQ, no stimulation of β-gal transcription was observed in the B3H experiment when the α-ProQ^ΔCTD^ fusion protein contained an R80A substitution (Fig 1E vs. D). The R80A variant is expressed at comparable levels to WT (Fig 1F), indicating that this loss of interaction is not due to destabilization of the fusion protein by the R80A mutation. We therefore concluded that use of a Δ*hfq* reporter strain allowed detection of specific ProQ-RNA interactions in our B3H assay, with comparable signal to what has been previously sufficient to conduct forward and reverse genetic analyses of Hfq RNA-binding surfaces.(30)

### Scope of detectable RNA interactions

Given that ProQ has been found to interact with dozens of sRNAs and hundreds of mRNAs inside of *Salmonella* and *E. coli* cells,(17) we sought to determine whether our B3H system could detect ProQ interactions with additional RNAs beyond *cspE*. We tested three additional RNAs that had been found to interact *in vivo* with ProQ: sRNAs SibB, RyjB and the 3’UTR of *fbaA* (hereafter, *fbaA*);(17, 22, 37) as well as four sRNAs for which we had previously detected B3H interactions with Hfq: ChiX, OxyS, ArcZ, and MgrR (Fig S3).(30) While a *cspE* hybrid RNA consistently produced the highest stimulation of transcription above basal levels when present with α-ProQ^Δ^CTD, we observed a ≥1.5-fold increase in fold-stimulation of β-gal activity when each of these hybrid RNAs was present in reporter cells (Fig 2A; full β-gal data in Fig S4A). As with *cspE*, β-gal activity arising from each of these hybrid RNAs was disrupted by an R80A point substitution (Fig 2B), suggesting the B3H signal represents a specific protein-RNA interaction. In addition, a hybrid RNA containing an arbitrary RNA sequence (the *trpA* terminator, T_trpA_) did not interact with the ProQ fusion protein. To compare the RNA-binding activity of ProQ to Hfq in this assay, we tested the same panel of nine hybrid RNAs for interaction with α-Hfq.(30) While interactions were detected for the four established Hfq-dependent sRNAs, α-Hfq did not stimulate β-gal transcription with any of the hybrid RNAs chosen as putative ProQ interactors (Fig 2C). We conclude that, against this panel of eight RNAs, and in the absence of endogenous Hfq, ProQ binds to a range of RNAs that co-immunoprecipitate with either ProQ or Hfq.(17, 38) As hybrid RNAs containing *cspE* and SibB yielded the highest B3H signal with ProQ, we focused further analysis on these two RNAs – one 3’UTR (*cspE*) and one sRNA (SibB).

**Figure 2.**
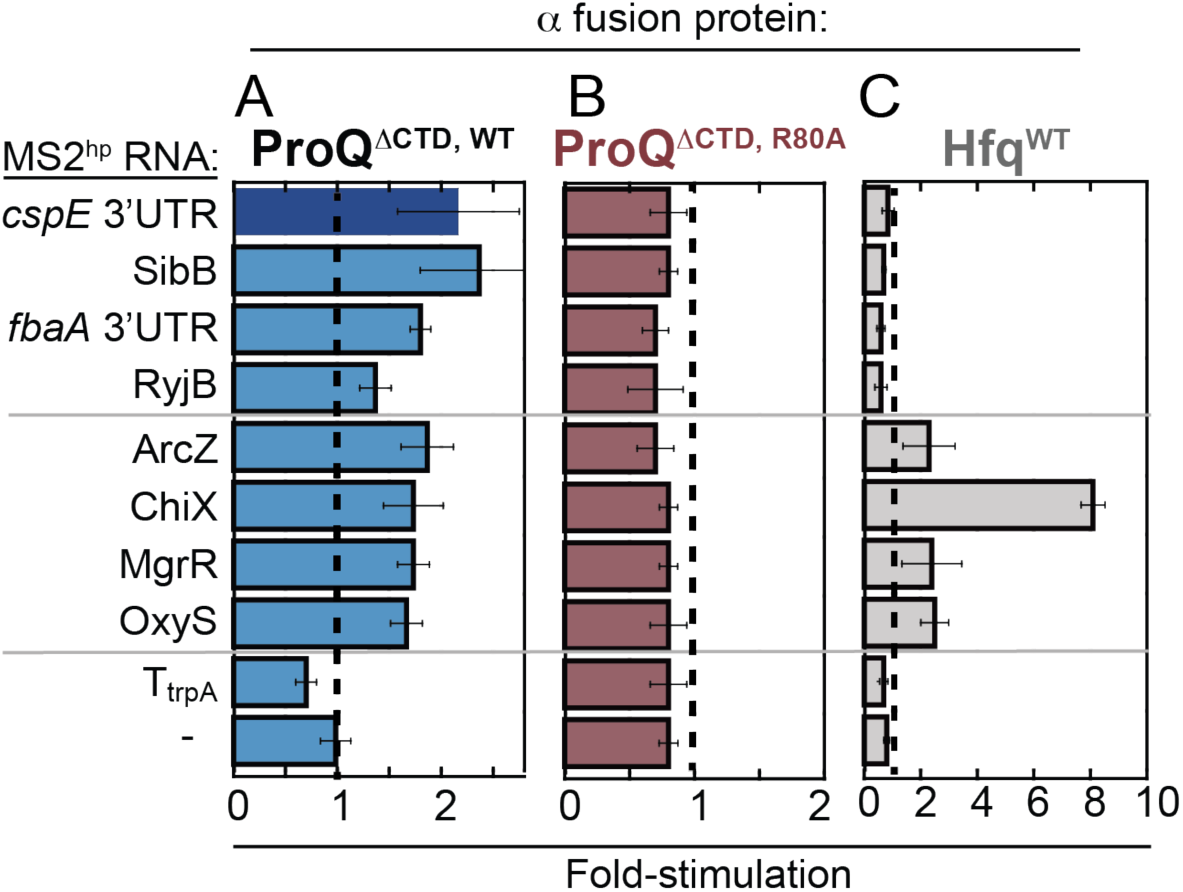
B3H assay detects ProQ’s interaction with multiple RNA substrates. Results of B3H assays between a panel of RNA substrates with (A) wild-type ProQ (B) an R80A variant or (C) wild-type *E. coli* Hfq. β-galactosidase assays were performed with *Δhfq* reporter strain cells containing three compatible plasmids: one that encoded λCI or the CI-MS2^CP^ fusion protein, another that encoded α or an α-fusion protein (α-ProQ^ΔCTD^, either with wild type ProQ or an R80A mutant, or α-Hfq), and a third that encoded a hybrid RNA (a single MS2^hp^ moiety fused to *cspE* 3’ UTR, SibB, *fbaA* 3’ UTR, RyjB, ArcZ, ChiX, MgrR, OxyS, trpA terminator (T_trpA_) or an RNA that contained only the MS2^hp^ moiety. The cells were grown in the presence of 0.2% arabinose and 50 μM IPTG (see Methods). The bar graph shows the fold-stimulation over basal levels as averages and standard deviations of values collected from three independent experiments conducted in triplicate across multiple days. Absolute β-gal values of a representative dataset are shown in Fig S4A.

### ProQ NTD is sufficient for binding *cspE* and SibB RNAs *in vivo*

Conflicting evidence has been collected about the contributions of the linker and CTD of ProQ to RNA binding.(20, 21, 28, 29) In order to assess the contribution of these ProQ domains to RNA binding *in vivo*, we compared the binding of five domain-truncation mutants of ProQ (Fig 3A; full β-gal data in Fig S4B). All fusion proteins were comparably expressed inside of the cell, as assessed by an antibody recognizing the region of α shared by each protein (Fig 3B). Removal of the CTD did not significantly alter the observed interactions with either *cspE* or SibB (FL vs. ΔCTD) and the CTD on its own did not afford any detectable interaction with either hybrid RNA (Fig 3C,D). Removal of the unstructured linker did not weaken ProQ’s interaction with *cspE* but did result in reduced interaction with SibB (ΔCTD vs. NTD). Interestingly, a construct with only the first 12aa of the 61-aa linker partially restored the interaction of ProQ with SibB (NTD+12aa vs. NTD; see Discussion). Together, our results indicate that the ProQ CTD is not required for interaction with either the *cspE* 3’UTR or SibB RNAs *in vivo* and that the NTD/FinO-domain is the primary RNA-binding site for both of these RNAs.

**Figure 3.**
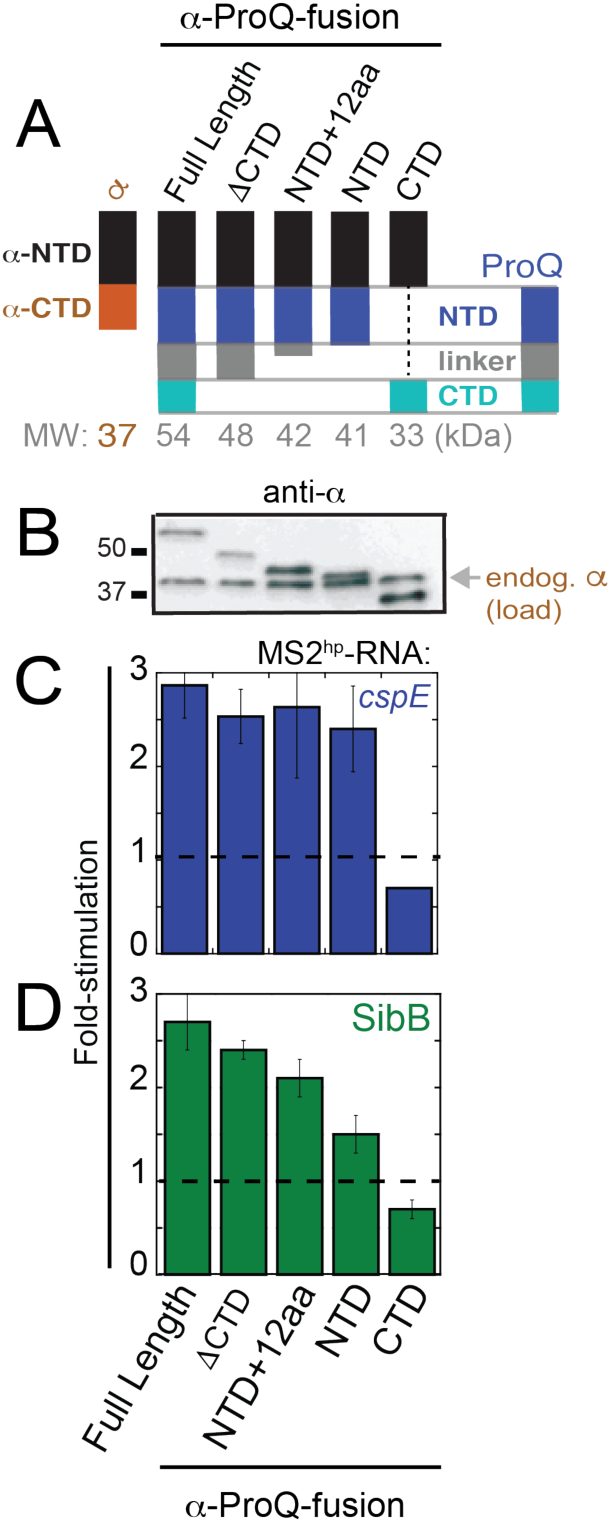
NTD is the primary site of interaction with *cspE* and SibB RNAs *in vivo*. (A) Schematic of α-ProQ domain-truncation mutants used in B3H assays, (B) Western blot with anti-RpoA antibody showing expression of α-ProQ truncations in lysates from samples in (C) and (D). The position of full-length endogenous RpoA (α; 37 kDa) and two molecular weight markers are indicated. Results of B3H assays detecting interactions between α-ProQ truncations and (C) *cspE* and (D) SibB RNAs. β-galactosidase assays were performed with *Δhfq* reporter strain cells containing three compatible plasmids: one that encoded λCI alone or the CI-MS2^CP^ fusion protein, another that encoded α or an α-fusion protein (α-ProQ^FL^ (full-length; resi=2-232), α-ProQ^ΔCTD^ (resi=2-176), α-ProQ^NTD+12aa^ (resi=2-131), α-ProQ^NTD^ (resi=2-119), or α-ProQ^CTD^ (resi=181-232)), and a third that encoded a hybrid RNA (MS2^hp^-*cspE* or MS2^hp^-SibB) or an RNA that contained only the MS2^hp^ moiety. The cells were grown in the presence of 0.2% arabinose and 50 μM IPTG (see Methods). The bar graph shows the fold-stimulation over basal levels as averages and standard deviations of values collected from three independent experiments conducted in triplicate across multiple days. Absolute β-gal values of a representative dataset are shown in Fig S4B.

### Conserved NTD residues mediate RNA interactions

Having established that the NTD/FinO-domain of ProQ is sufficient for interaction with both *cspE* and SibB RNAs *in vivo*, we wanted to identify amino acids in the NTD beyond Arg80 that are required for RNA interaction. Hereafter, we call the two faces of the ProQ NTD the “concave face” (containing H2 and H3 as primary structural features) and “convex face” (containing H1 and β1/2 as structural feature; Fig S1A, Fig 4A) to be consistent with nomenclature used for other FinO-domain proteins.(16) Mapping degree-of-conservation onto the ProQ NMR structure, we noticed a large patch of highly conserved residues on the opposite face as Arg80 (Fig S1B), and wondered whether these conserved “concave-face” residues are important for RNA binding. To explore this possibility, we identified residues that are both highly conserved across 15 ProQ/FinO-domain proteins (Fig S5) and surface exposed in the NMR structure to target for mutagenesis in the α-ProQ^ΔCTD^ construct (Fig 4A-C).

**Figure 4.**
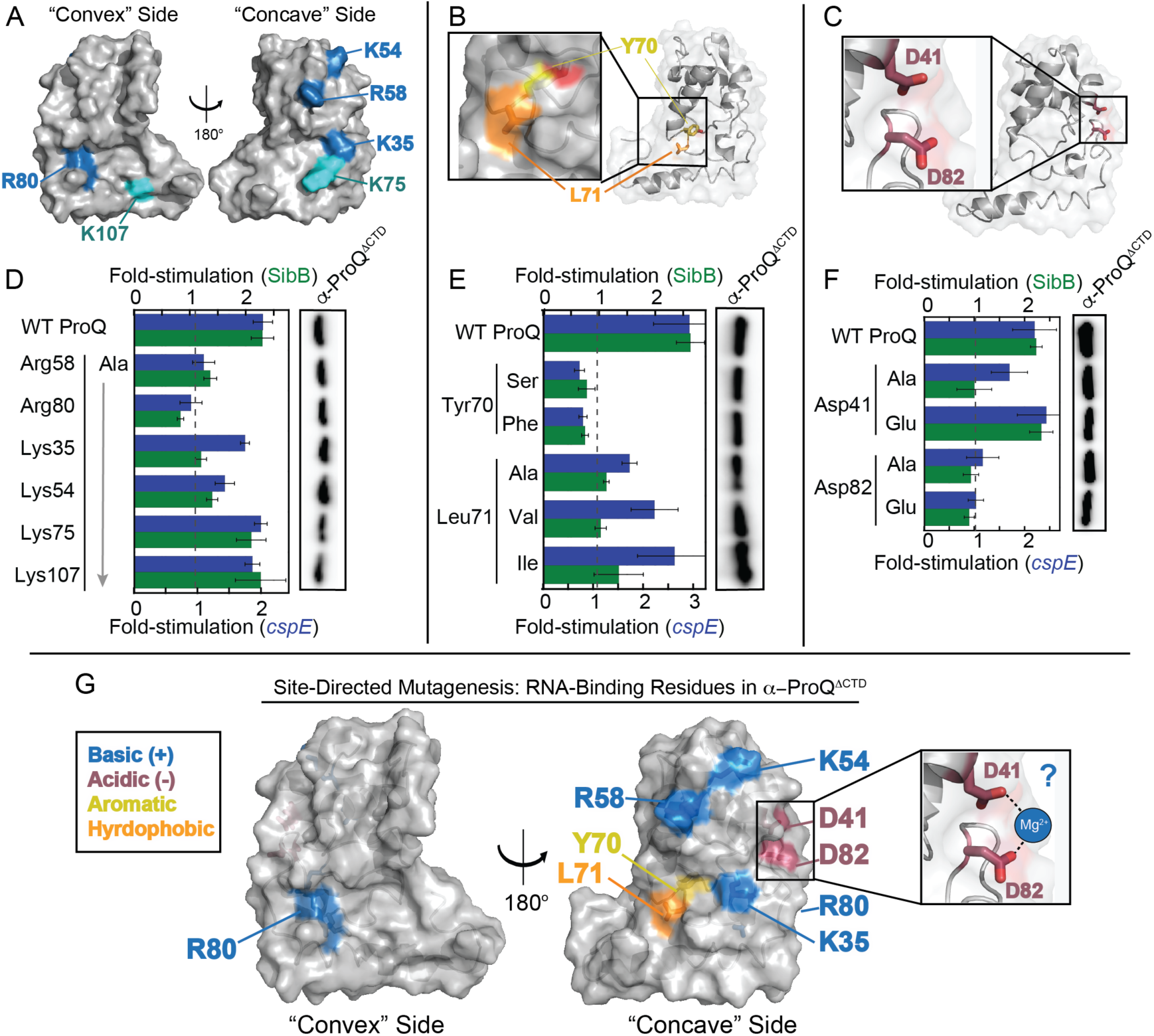
Effects of specific disruptive amino acid substitutions on B3H interactions. Positions of (A) basic, (B) hydrophobic and (C) acidic residues targeted for site-directed mutagenesis shown on ProQ NTD Structure (PDB ID: 5nb9).(28) Residue coloring, used throughout: highly conserved basic, blue; less conserved basic, cyan; hydrophobic, orange; aromatic, yellow; acidic, red). (D-F, left) Results of B3H assays showing effects on ProQ-RNA interactions of alanine mutations at (D) basic, (E) hydrophobic and (F) acidic residues. β-galactosidase assays were performed with *Δhfq* reporter strain cells containing three compatible plasmids: one that encoded λCI or the CI-MS2^CP^ fusion protein, another that encoded α or an α-ProQ^ΔCTD^ fusion protein (wild type, WT, or the indicated mutant), and a third that encoded a hybrid RNA (MS2^hp^-*cspE* or MS2^hp^-SibB) or an RNA that contained only the MS2^hp^ moiety. The cells were grown in the presence of 0.2% arabinose and 50 μM IPTG (see Methods). The bar graph shows the fold-stimulation over basal levels as averages and standard deviations of values collected from three independent experiments conducted in triplicate across multiple days. Absolute β-gal values of a representative dataset are shown in Fig S6. (D-F, right) Western blot to compare steady-state expression levels of mutant α-ProQ^ΔCTD^ fusion proteins. Lysates were taken from the corresponding β-gal experimeent containing MS2^hp^-*cspE* and all other hybrid components at 50 μM. Following electrophoresis and transfer, membranes were probed with anti-ProQ antibody (see Fig S2). (G) Summary of results from site-mutagenesis experiments. Surface representation of ProQ NTD structure (PDB ID: 5nb9)(28) viewed from convex face (left) and concave face (right). Residues at which substitution with alanine disrupts RNA binding are colored as above. Inset on right shows ProQ structure as a cartoon under a transparent surface with side chains of Asp residues as sticks and a putative magnesium ion (Mg^2+^) is shown in blue coordinating the Asp residues. This Mg^2+^ was not modeled in the NMR structure, but we speculate it may be involved in RNA binding (see Discussion).

Given the negative electrostatic nature of RNA, we focused first on contributions of positively charged residues on the NTD surface. The recent NMR structure of *E. coli* ProQ revealed that both faces of the NTD/FinO-domain possess patches of positively charged residues (Fig S1C),(28) leading to ambiguity about which would be most important for RNA binding. We selected six basic residues to substitute individually with alanine – four highly conserved (Lys35, Lys54, Arg58, Arg80; Fig 4A, blue) and two more modestly conserved (Lys75, Lys107; Fig 4A, cyan). Altered forms of the ProQ fusion protein were expressed at levels comparable to WT (Fig 4D, right) and removal of highly conserved basic residues (R58A, R80A, and K54A variants) strongly reduced interaction with *cspE* and SibB hybrid RNAs, while the K35A variant demonstrated preferential loss of interaction with SibB (Fig 4D; full β-gal data in Fig S6). Substitution of the less conserved basic residues with alanine (K75A and K107A variants) had more modest effects on RNA interaction (Fig 4D). Together, these results suggest that the concave face of the NTD/FinO-domain contributes to RNA binding along with Arg80 on the convex face, and that conservation of surface-exposed residues correlates with their role in RNA binding.

Two of the most highly conserved residues along the NTD’s concave face are aromatic and hydrophobic residues in which the side chains are partially surface exposed: Tyr70 and Leu71 (Fig 4B). As such residues can mediate intermolecular interactions, we wished to determine whether they contribute to RNA binding. Substitution of Leu70 with alanine significantly reduced the interaction of ProQ^ΔCTD^ with both *cspE* and SibB hybrid RNAs without affecting expression levels of the ProQ fusion protein (Fig 4E). Even very conservative substitutions at this position (Ile or Val) resulted in decreased RNA interaction with both hybrid RNAs (Fig 4E), consistent with hydrophobic interactions depending on the size and shape of aliphatic chains. When the neighboring Tyr70 residue was altered, both Y70S and Y70F ProQ variants showed a loss of RNA interaction despite WT-levels of expression (Fig 4E). That Phe and Ser are each insufficient for RNA interaction at this position indicates that both the hydroxyl group and aromatic ring of Tyr70 are critical for RNA interaction with *cspE* and SibB (see Discussion).

We next examined the role of two highly conserved acidic residues, Asp41 and Asp82, positioned only ∼5 Å apart in the folded protein (Fig 4C). Alanine substitution at each position strongly impaired interaction of ProQ with SibB, while interaction with *cspE* was less impacted by D41A than a D82A substitution (Fig 4F). We next tested subtler structural changes at these positions by replacing Asp with Glu, extending the side chain by a single –CH_2_– group. While α-ProQ^ΔCTD^ with a D41E substitution was fully able to interact with both *cspE* and SibB hybrid RNAs, a D82E substitution strongly impaired interaction with both RNAs (Fig 4F). Importantly, all four ProQ variants were comparably stable to WT (Fig 4F, right). Together, these results suggest that a pair of Asp residues, along with the precise positioning of the Asp82-carboxylate moiety, is critical for RNA interaction. These negatively charged aspartate residues could contribute to RNA binding either through hydrogen bonding or through coordination of a magnesium ion (Fig 4G, inset; see Discussion). Together, our site-directed-mutagenesis results demonstrate the importance of numerous residues – basic, acidic and hydrophobic – across the conserved concave-face of the ProQ NTD, along with Arg80 on the opposite surface, in mediating RNA interactions with *cspE* and SibB (Fig 4G).

### Unbiased genetic screen confirms role of concave face in RNA interactions

Our site-directed-mutagenesis results strongly implicated the concave face of the NTD/FinO-domain as a critical site for RNA binding in the context of ProQ^ΔCTD^, but it is possible this analysis overlooked other critical regions of the protein. We therefore used our genetic B3H assay to conduct an unbiased forward genetic screen to identify ProQ residues critical for RNA binding. We began with a library of mutagenized plasmids containing full-length *proQ* (α*-proQ*^*FL*^) to leave open the possibility of finding substitutions anywhere in the protein that would disrupt interaction with either *cspE* or SibB hybrid RNAs. For the screen, B3H reporter-strain cells containing the CI-MS2^CP^ adapter protein and either the MS2^hp^-*cspE* or MS2^hp^-SibB hybrid RNA were transformed with a PCR-mutagenized α-*proQ*^*FL*^ plasmid library estimated to contain ∼23,000 unique mutants, and plated on X-gal indicator medium (see Methods). In this primary screen, ∼15% of colonies were white or pale, the phenotype expected for transformants that contained α-*proQ* mutants that no longer interacted with a hybrid RNA (Fig S7A). To eliminate the subset of colonies containing plasmids encoding mutations resulting in unstable fusion proteins, we established a dot-blot assay in which lysates from single colonies could be spotted on nitrocellulose membranes and probed with an anti-ProQ antibody. Indeed, the dot-blot assay displayed strong signal above an α-empty negative control, even in a *proQ*^*+*^ reporter strain, and a suitable linear range for the intended counter-assay (Fig S7B; see Methods). From the 536 white or pale colonies identified in the primary screen (372 isolated against *cspE* + 164 against SibB RNA), the dot-blot assay identified the subset (∼30%) that maintained wild-type levels of expression (Fig S7C).

We sequenced α-*proQ* plasmids from colonies displaying strong defects in RNA binding while retaining high levels of fusion-protein expression (Fig S7D, purple oval). Sequencing reads unambiguously covering the entirety of the *proQ* sequence were obtained for 86 mutant plasmids, of which 54 were found to harbor a single mutation; nearly a third of mutant plasmids were independently isolated multiple times (Table 1). Together, these plasmids encoded 37 distinct amino-acid substitutions at 25 residues in ProQ that disrupt interaction with one or both RNAs used in our screen (Table 1). We confirmed the loss of RNA interaction of these 37 α-ProQ^FL^ variants in liquid β-gal assays with both *cspE* and SibB hybrid RNAs, and verified their stability via dot-blot assays (Table S5). These experiments were conducted at two IPTG concentrations to examine RNA-binding across a range of α-ProQ^FL^ expression levels. Results from these experiments demonstrate that none of the RNA-binding defects of the 37 α-ProQ^FL^ variants identified here are attributable to reduced protein expression relative to WT.

**Table 1.** Results of forward genetic screen for ProQ substitutions that disrupt RNA binding. Each row represents a plasmid isolated from the screen one or more times which expressed a variant α-ProQ^FL^ protein that was expressed at wild-type or greater levels and nevertheless displayed reduced β-galactosidase activity with either *cspE* or SibB hybrid RNAs. Columns indicate (1) the residues in α-ProQ^FL^ at which substitutions were found to disrupt RNA binding in B3H screen, (2) the position of each residue based on the ProQ NTD NMR structure (PDB: 5nb9),(28) either within the core of the protein, on the surface or buried, but on the periphery outside of the core (see Fig S8), (3) the specific amino-acid substitution resulting from mutation in each isolated plasmid and (4) the number of times this mutated plasmid was isolated in screening against either a MS2^hp^-*cspE* or MS2^hp^-SibB RNAs.

| ProQ Residue | Location in NTD | Substitution | Times Isolated |  |
| --- | --- | --- | --- | --- |
|  |  |  | w/ *cspE* | w/ SibB |
| L17 | core | L17P | 1 | 1 |
| R20 | surface | R20P |  | 1 |
| F21 | buried | F21S |  | 1 |
| C24 | core | C24W | 1 |  |
|  |  | C24R |  | 1 |
| F25 | core | F25C | 1 |  |
|  |  | F25S | 2 | 1 |
|  |  | F25Y |  | 1 |
| L34 | core | L34R | 1 |  |
|  |  | L34Q |  | 1 |
|  |  | L34P |  | 1 |
| K35 | surface | K35E |  | 1 |
|  |  | K35N |  | 1 |
|  |  | K35I |  | 1 |
| G37 | surface | G37V |  | 1 |
| I38 | core | I38S | 1 |  |
| L42 | core | L42S |  | 1 |
| L57 | core | L57S | 1 |  |
| A60 | core | A60D | 1 |  |
| L63 | surface | L63P |  | 1 |
| Y64 | core | Y64C | 1 |  |
|  |  | Y64N | 1 | 1 |
| S66 | surface | S66P | 1 | 1 |
| Y70 | surface | Y70H | 2 | 1 |
| L71 | surface | L71P | 2 | 1 |
| R80 | surface | R80C | 1 |  |
|  |  | R80H | 1 |  |
|  |  | R80S |  | 2 |
| V81 | core | V81D | 2 | 1 |
| D82 | surface | D82Y | 1 |  |
| L83 | core | L83F | 1 |  |
|  |  | L83P |  | 1 |
| G85 | surface | G85D | 1 | 1 |
| L91 | buried | L91Q |  | 1 |
|  |  | L91R | 1 |  |
| Q102 | surface | Q102P |  | 1 |
| L103 | buried | L103P |  | 1 |
| **25 residues** |  | **37 variants** | **24 plasmids** | **26 plasmids** |

Notably, despite beginning this screen with a library of mutations in full-length *proQ*, all 25 residues implicated by the screen in RNA binding are located in the FinO-like NTD. Nearly all ProQ variants, whether identified in the screen against either RNA, resulted in diminished interaction with *both cspE* and SibB hybrid RNAs (Table S5), suggesting that ProQ binds both of these RNAs with a similar surface and molecular mechanism (see Discussion). Of the implicated residues, the NMR structure shows 14 are buried in the protein structure, while 11 residues are surface-exposed (Fig S8A-C).(28) To validate the screen’s results, we set aside variants at buried residues as likely to perturb the overall structure of the protein, and further set aside surface-exposed residues we had already investigated through site-directed mutagenesis (Lys35, Tyr70, Leu71, Arg80, Asp82). This left six previously unexamined surface-exposed residues suggested by our screen to contribute to RNA binding (Fig S8E). Many mutations identified by the screen at these positions produced non-conservative substitutions, such as the introduction of a proline residue (Table 1; Fig S8C). To determine whether loss of RNA interaction for each variant arose from the absence of a wild-type residue or the presence a destabilizing one, we made site-directed alanine substitutions in α-ProQ^ΔCTD^ at each position. When Arg20, Leu63, Ser66 and Gln102 were each replaced with alanine, RNA binding was not strongly impaired (Fig S8E,F). Substitutions at these positions with proline likely emerged from our screen due to structural disruption by proline rather than the native residues contributing essential molecular contacts with RNA. In contrast, alanine substitutions at two glycine positions (Gly37 and Gly85) strongly disrupted RNA binding without affecting expression of each fusion protein (Fig S8D-F). We conclude that these three residues are critical for the NTD’s interaction with RNA. This allowed us to map on to the ProQ NMR structure all of the validated residues identified by our forward-genetic screen to be necessary for RNA interaction (Fig 5A). This highlights a patch of RNA-binding residues along the concave face and wrapping around to Arg80. In addition to residues already probed through site directed-mutagenesis, Gly37 is located on the conserved concave face of the NTD (Fig 5A), while Gly85 is a part of the β3-4 hairpin that contains Arg80 and Asp82 (Fig 5B; see Discussion).

**Figure 5.**
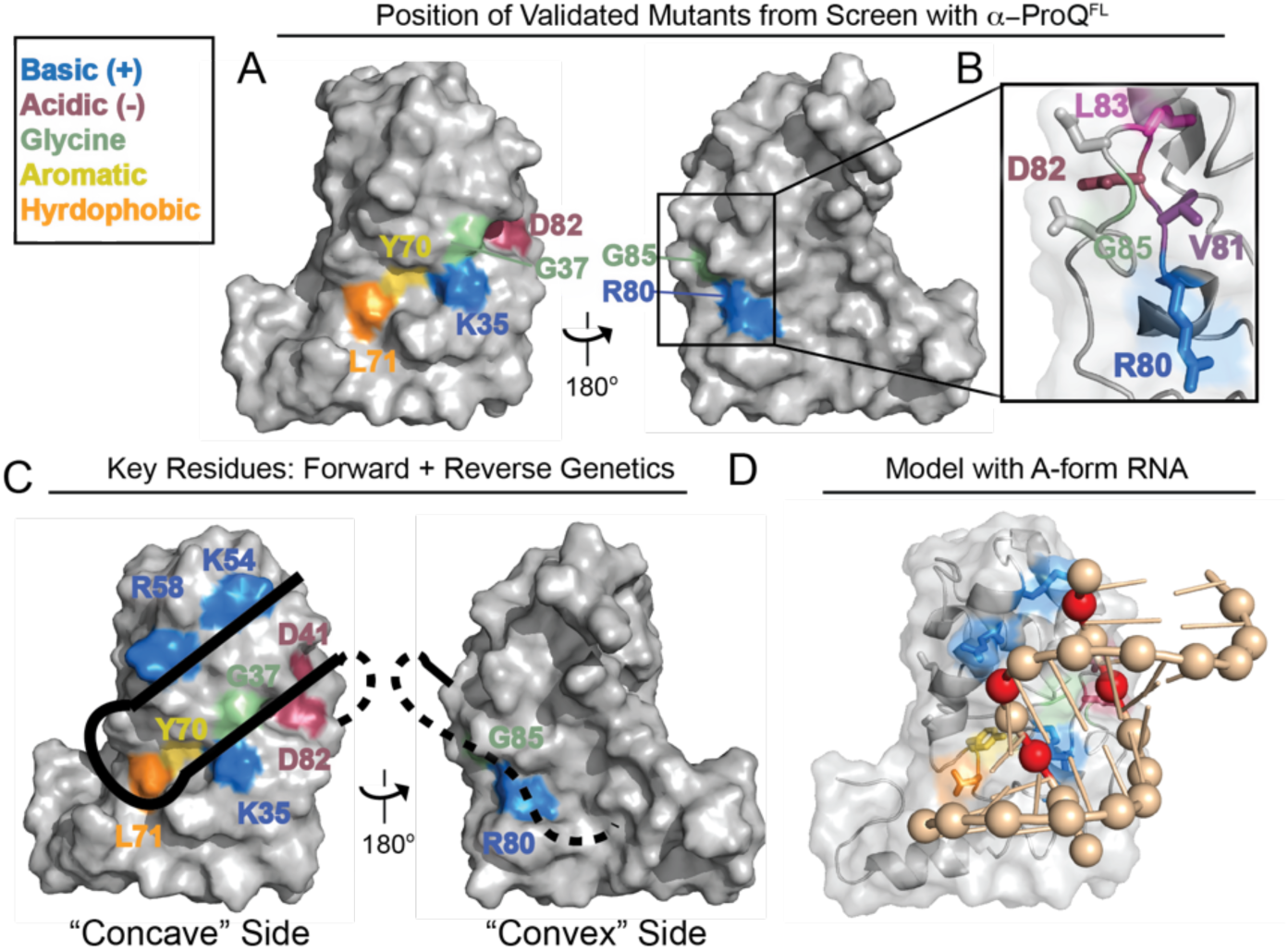
Validated genetic-screen results and model for ProQ-dsRNA interactions. (A) Surface representation of ProQ NTD structure (PDB ID: 5nb9),(28) viewed from concave (left) or convex (right) surface, showing residues at which substitutions were found to disrupt RNA binding in B3H screens of mutagenized α*-ProQ*^*FL*^ plasmids and at which substitution with alanine has been confirmed to be sufficient to disrupt binding (Fig 4 and Fig S8E; residue coloring: basic, blue; hydrophobic, orange; aromatic, yellow; acidic, red; polar: purple; glycine: green) (B) Inset shows close-up view of β3/4 hairpin, viewed from the convex face. Under a transparent surface representation, the polypeptide backbone is shown as a cartoon and amino-acid side chains are represented sticks, colored as in (A). Side chains for residues Asp84 and Asn86 (not identified as RNA-binding residues in screen) are shown as grey sticks. (C) Summary of results from both site-directed and unbiased mutagenesis experiments. Surface representation of ProQ NTD structure, viewed from concave (left) or convex (right) surface, showing all residues identified in this study as necessary for strong RNA interactions *in vivo* whether from site-directed mutagenesis (Fig 4) or a forward genetic screen (A) and colored as in (A). A schematic cartoon of an RNA hairpin is shown over the concave face, representing one mode of binding that would be consistent with these results. (D) Preliminary structural model for ProQ NTD recognition of dsRNA. Transparent surface and cartoon representation of ProQ NTD viewed from concave face, hand-docked in PyMol (version 1.6.2) to a 12-bp RNA duplex (PDB ID: 5DA6),(47) using only rigid rotations of the protein and RNA structure. RNA as a tan cartoon with phosphates as spheres. Four phosphates that have a suitable geometry to interact with basic and acidic residues (Lys35, Lys54, Arg58 and Asp41, Asp82) are colored in red. ProQ and residues are colored as in (C) with side chains of RNA-binding residues shown as sticks.

### Potential for dsRNA recognition by ProQ RNA-binding residues

While there are multiple ways that the ProQ residues we have identified could contribute to RNA binding, we wished to develop a preliminary structural model that would ∂combine results from our forward and reverse genetic approaches (Fig 5C) with literature results suggesting a strong preference for ProQ to bind structured RNAs,(22) and FinO requiring both a stem and neighboring ssRNA for strong binding to FinP RNA.(26, 27) We noticed that the group of RNA-binding residues we have identified spans 15-20 Å across the concave face of the NTD, similar to the width of an A-form RNA helix. Thus, we propose a model in which the concave face of the NTD/FinO-domain recognizes the duplex region of an RNA substrate, with Arg80 on the convex face potentially interacting with a more flexible region of nearby RNA (Fig 5C). As electrostatics often predominate interaction with RNAs, we examined the positions of three conserved basic concave-face residues (Arg58, Lys54, Lys35) and two conserved acidic residues (Asp41, Asp82) implicated in RNA binding by our genetic analyses. Docking of a duplex RNA structure onto the ProQ NTD structure shows that these five residues are positioned in such a way to facilitate electrostatic interactions with phosphates within an A-form-RNA helix (Fig 5D; phosphates proposed to be contacted = red spheres). It is also possible that Asp41 and/or Asp82 contribute H-bonding interactions directly to the RNA. Leu71 is positioned on the other end of the electrostatically charged patch on the concave surface and could mediate interactions with the hydrophobic face of a nucleobase in a nearby RNA element, such as the loop of a hairpin. Based on this preliminary modeling, we propose that the ProQ NTD/FinO-domain could serve as a scaffold for the patterned display of charged residues, positioned in such a way to recognize the shape of the negatively-charged backbone of duplex RNA.

## DISCUSSION

In this work, we have conducted the first comprehensive mutagenesis study of ProQ to identify the functional surface used by ProQ to bind to RNA substrates. In order to apply both forward and reverse genetic approaches to this question, we adapted a bacterial three-hybrid (B3H) assay for genetic detection of RNA-protein interactions to report on binding of ProQ to RNA. Using this system, we have established that the NTD/FinO-domain of ProQ is the primary site that mediates interaction with *cspE* and SibB RNAs *in vivo* and have dissected the roles of residues on multiple surfaces of this domain in RNA binding. Results from both forward- and reverse-genetic analyses are in strong agreement with one another, converging to implicate the more highly conserved surface of the NTD, which we call the concave face, as the primary site for recognition of both RNAs. We have proposed a working structural model that interprets the positions of residues identified in this study as critical for RNA binding in light of ProQ’s established preference for binding to structured RNAs. In this model, the global structure of the NTD/FinO-domain pre-positions highly conserved residues across the concave face to recognize the conserved shape and charge of double-stranded RNA. By allowing conservation to guide our initial studies and taking an unbiased genetic approach, we have demonstrated that, along with basic residues, conserved acidic, aromatic and hydrophobic residues also play an important role in RNA binding by ProQ.

### Insights into RNA-binding surfaces of ProQ

The relative roles in RNA binding played by disordered regions and structured domains of ProQ/FinO-like proteins have been a subject of inquiry over several years.(20, 21, 28, 29) Two lines of evidence from our study suggest that the ProQ NTD/FinO-domain is the primary *in vivo* binding site for interaction with the two RNAs closely investigated here. First, truncation analysis demonstrates that the ProQ NTD, along with the first 12 aa of the linker, is sufficient for full interaction with both the 3’ UTR of *cspE* and the sRNA SibB. Further, an unbiased forward genetic screen starting with mutagenized full-length *proQ* did not identify any mutations in the region encoding the linker or CTD that were sufficient to disrupt interaction of ProQ with either hybrid RNA. In contrast, 37 amino-acid substitutions at 25 residues within the NTD/FinO-domain were identified by our screen to disrupt interactions with one or both hybrid RNAs.

While our data support a model in which the NTD/FinO-domain of ProQ is the primary binding site for these two RNAs *in vivo*, we cannot rule out the possibility that other regions in the linker or CTD contribute important interactions with certain RNA substrates. In particular, our data suggest that the most N-terminal 12 aa of the unstructured linker may be necessary for full interaction with certain RNA substrates (*e.g.* SibB), while not for others (*e.g. cspE*; Fig 3). Among these 12 aa in *E. coli* ProQ are four basic residues, three of which are immediately adjacent to one another (Fig S5), and the presence of positively charged residues in this region are common across other ProQ proteins. It is possible that certain RNAs may depend on electrostatic stabilization from the NTD-adjacent region of the linker. It will be interesting to explore this possibility in the future with our panel of RNA substrates using both forward- and reverse-genetic approaches.

Within the NTD/FinO-domain, the majority of RNA-binding residues identified in this study map to the concave face, but Arg80 on the convex face of ProQ is essential for interaction with all eight RNAs we have tested (Fig 2). The observation that positively charged residues on both surfaces of the NTD/FinO-domain contribute to RNA binding is consistent with crosslinking studies with *E. coli* FinO that found basic residues on both faces crosslink to FinP RNA, and with biophysical studies suggesting RNA binding on the convex face of ProQ.(28, 29) It is striking, however, that Arg80 is the only residue on the convex face of the NTD that our unbiased genetic screen implicated in RNA binding. One intriguing possibility is that, while the concave face may mediate interactions with duplex RNA, the convex face may interact with nearby single-stranded region(s) (Fig 5C). It will be important to explore the mechanistic role of Arg80 in RNA-binding further in the future.

### Conservation and connections with other FinO-domain proteins

Many of our findings align well with previous results obtained with ProQ and other FinO-domain proteins. For instance, the critical role of the NTD/FinO-domain in RNA interactions is consistent with *in vitro* findings that the FinO-domain of *E. coli* ProQ, and also of *L. pneumophila* RocC, is sufficient for high affinity binding to its RNA substrates.(20, 21) Further, many of the concave-face RNA-binding residues we have identified in *Ec* ProQ are conserved in both of these homologs, as well as in the FinO-containing *N. meningitidis* NMB1681 (Fig S9). Previously determined crystal structures of NMB1681 and F’ FinO reveal that these conserved residues also map largely to the concave faces of their respective FinO-domain (Fig S9B-D).(39, 40) In addition, the two positions in FinO that crosslink most strongly to FinP RNA in previous work are located on helix H3, in similar positions to Lys54 and Arg58 on the concave face of *Ec* ProQ.(29)

A universally conserved residue across all of these FinO-domain proteins is the aromatic residue Tyr70 (*Ec* numbering; Fig S9A), which appears to play a critical role in the structure and/or function of ProQ. In our random-mutagenesis screen, a Y70H substitution was identified independently as disrupting *cspE* and SibB interactions. Interestingly, a Y-to-F mutation at the analogous position was found in an unbiased screen to disrupt RocR activity in *L. pneumophila*,(20) and the same Y70F substitution in *E. coli* ProQ impairs binding with both RNAs we have examined, reaffirming the importance of this hydroxyl group. In the ProQ NMR structure, the aromatic ring of Tyr70 is somewhat buried while the hydroxyl group is pointing towards the surface (Fig 4B),(28) and we cannot rule out that the role of Tyr70 in RNA binding may be mediated at least partially through global structure of ProQ. We note this hydroxyl group is relatively close to backbone amides of Leu34 and Lys35 in *Ec* ProQ (2.5-4.7 Å in various NMR states) and could mediate an intramolecular hydrogen bond within the polypeptide,(28) or could be directly involved in contacting RNA.

Two glycine residues (Gly37 and Gly85) were found in our unbiased screen to be necessary for RNA interaction; even an alanine at these positions prevents interaction with *cspE* and SibB RNAs (Fig S8E). Both glycine residues are highly conserved in other FinO-domain proteins (Fig S9) and located near structural elements that contain additional RNA-binding residues: at the base of H3 between Lys35 and Asp41, and in the β3-4 hairpin containing Arg80 and Asp82 (Fig 5B). One interpretation of these results is that a flexible polypeptide conformation at these sites is critical to facilitate RNA binding. The structural role of Gly85 is especially interesting as our data implicate this β3-4 hairpin as a critical structural element for RNA interaction by ProQ. While this structural feature is also found in structures of *N. meningitides* NMB1681 and *E. coli* F’ FinO (Fig S9C,D),(28, 39, 40) this region in FinO does not crosslink to FinP RNA (29) and had not been previously appreciated as an element contributing to RNA binding. In this study, however, the β3-4 hairpin featured the highest density of hits in our unbiased genetic screen: in addition to Arg80, Asp82, and Gly85, two additional hydrophobic residues in this β3-4 hairpin (Val81 and Leu83) were disrupted by mutants isolated in our screen. The latter residues appear to pack the β3-4 hairpin into the global core of the ProQ NTD (Fig 5B) and are part of a large number of “core” residues identified by our screen at which substitutions disrupt RNA binding without affecting protein expression levels (Fig S8A,B; Table S5).

### Subtle substitutions with dramatic RNA-binding effects

A striking feature of our results is the large number of minor chemical perturbations that nevertheless strongly impact ProQ’s interaction with RNA. In addition to substitutions at many “core” residues that were shown by our screen to disrupt RNA binding without affecting ProQ expression levels (Fig S8A-B), we found that subtle variations of surface amino-acids have striking effects on RNA binding. Remarkably, even a slight perturbation in the positioning of the Asp82-carboxylate group arising from a D-to-E substitution is sufficient to severely impair RNA binding (Fig 4F). We envision that Asp82 – present in the β3-4 hairpin that also contains Arg80 and Gly85 – contributes to RNA binding either through hydrogen bonds to an RNA nucleobase or by coordinating a metal ion together with nearby Asp41, which is especially important for SibB interaction (Fig 4F). In the latter case, an Asp-bound cation such as magnesium (Mg^2+^; Fig 4G, inset) could serve as a fourth positively-charged moiety spanning the conserved patch of the NTD’s concave face, and could potentially interact with a phosphate, as we have preliminarily modeled (Fig 5D). Subtle substitutions of Leu70 with alternate aliphatic residues also have surprisingly strong effects on RNA binding, with even isoleucine – a structural isomer of leucine – impairing SibB binding (Fig 4E). Considered together with contributions to RNA binding by residues spanning a wide area across the concave face of the NTD, the disruptive effects of subtle substitutions in both the core and on the surface of ProQ suggest that the global structure of the ProQ NTD/FinO-domain mediates RNA recognition through precise positioning of multiple chemical moieties at a specific distance and orientation to one another. Our preliminary model of RNA recognition proposes that the precise positioning of these chemical groups acts to read out the shape and charge of an RNA duplex, consistent with evidence suggesting ProQ selects RNA substrates based as much on structure as sequence.(22)

### Relationship between ProQ and Hfq

There has been interest in the overlap of the subset of cellular RNAs bound by ProQ and Hfq,(22, 37) another global RNA-binding protein that stabilizes dozens of sRNAs and catalyzes annealing with mRNAs in *E. coli*. In this study, ProQ was found to bind to a wider range of RNAs than Hfq, producing B3H interactions with RNAs found to interact both with ProQ as well as with Hfq *in vivo* (Fig 2). Deletion of endogenous *hfq* from the *E. coli* reporter strain resulted in a strengthened B3H interaction of ProQ and RNA, even more so than deletion of endogenous *proQ* (Fig 1). While it is tempting to speculate that this reflects competition of ProQ with Hfq for RNA substrates, it is notable that we do not observe B3H interactions of α-Hfq with *cspE*, SibB, *fbaA* and RyjB, suggesting that any interaction between Hfq and these RNAs is likely weak relative to Hfq-dependent sRNAs. An *Δhfq* reporter strain was previously found to be ideal for Hfq-sRNA B3H interactions.(30) While it is possible that this strain benefits both Hfq- and ProQ-RNA interactions by eliminating competition between endogenous Hfq and the RNAP-bound fusion protein, it is also possible that the benefit arises due to a pleiotropic, indirect effect of *Δhfq*.(41) Collectively, our data are consistent with ProQ and Hfq sharing a subset of RNA targets, as has been suggested by previous studies.(22, 37)

It is well established that Hfq has multiple RNA-binding surfaces that possess distinct RNA-binding specificity and contribute to RNA annealing.(6, 42) While *E. coli* FinO and *N. meningitidis* NMB1681 have been shown to catalyze RNA annealing and strand exchange,(18, 19, 21, 40) these activities have not yet been demonstrated for ProQ itself. Based on analogy to Hfq function, a likely prerequisite for RNA annealing would be the ability to bind multiple RNA substrates simultaneously on distinct binding surfaces. Here, we have investigated the domains and surfaces that mediate interaction with one sRNA and one mRNA 3’UTR. SibB is a *cis*-encoded antitoxin sRNAs and thus may possess more extensive complementarity with its cognate toxin mRNAs than most Hfq-dependent sRNAs.(43, 44) The vast majority of mutations we have examined here have strikingly similar effects on the binding of *cspE* and SibB hybrid RNAs, with a few intriguing exceptions. For instance, the interaction of ProQ with SibB in our B3H assay depends more on the ProQ linker, and on residues Lys35 and Asp41, than that with *cspE*. We look forward to searching for additional RNA-specific binding effects of *proQ* mutations in the future, using a larger set of interacting RNAs, and determining to what extent SibB and *cspE* represent apparent RNA “classes” of sRNAs and 3’UTR, as well as exploring interactions of 5’UTR-fragments and coding regions, which recent datasets show are quite abundant in ProQ-bound pairs of RNAs.(37)

### Outlook

Many questions remain about the structure and function of ProQ, including *(i)* what the detailed role of the Arg80-containing convex face in RNA binding, *(ii)* to what extent unique modes of interaction exist for distinct RNAs or classes of RNAs, *(iii)* whether ProQ mediates RNA annealing and which part(s) of ProQ would contribute to this activity, *(iv)* which part(s) of ProQ may recruit additional cellular factors, such as the ribosome,(45) RNA polymerase, PNPase (46) or other factors. The genetic assay we have developed could be useful in several of these pursuits: genetic screens conducted with counter-screens against various RNAs have the potential to identify ProQ substitutions with RNA-specific binding effects. The fact that our interaction assay is conducted *in vivo* means that interactions we detect could be influenced by one or more of the above cellular factors. It is intriguing to imagine that a chromosomal screen could be used to identify cellular factors that influence the state of ProQ-RNA interactions. In addition, the ProQ variants identified in this work will serve as helpful tools to probe the contribution of RNA binding by distinct surfaces to cellular pathways of gene expression. Finally, we look forward to comparing our preliminary genetically-guided model for ProQ’s interaction with duplex RNA with a high-resolution co-structure of this complex. Indeed, we anticipate that the model presented in this study can guide future strategies to obtain such a high-resolution structure. The structural details of this protein-RNA recognition event provide an important foundation to further elucidate molecular mechanisms of gene regulation by the global RNA-binding protein ProQ.

## SUPPLEMENTARY DATA

Supplementary Data are available as a PDF online.

## AUTHOR CONTRIBUTIONS

S.P., C.M.G., and K.E.B. conceived the ideas and designed experiments. S.P., C.M.G., and O.M.S. performed B3H experiments. S.P. and C.M.G. performed the genetic screen. C.M.G. performed immunoblotting experiments. S.P., C.M.G., O.M.S., C.D.W., C.L.H., H.L. and K.E.B. performed molecular cloning to generate key resources. S.P. and K.E.B. wrote the original draft of the manuscript. S.P., C.M.G., O.M.S., C.D.W., C.L.H., H.L. and K.E.B. contributed toward review and editing of the manuscript.

## ACKNOWLEDGEMENTS

We thank members of the Berry, Camp and Lijek laboratories for comments on the manuscript, advice and discussion. We thank Gizela Storz for generously providing an aliquot of an anti-ProQ antibody and also for helpful comments and discussion about the manuscript.

