## Supplementary_Data for "Genetic identification of the functional surface for RNA binding by *Escherichia coli* ProQ"

This pdf file includes:

Supplementary Figures S1 to S9

Supplementary Tables S1 to S5

Supplementary References

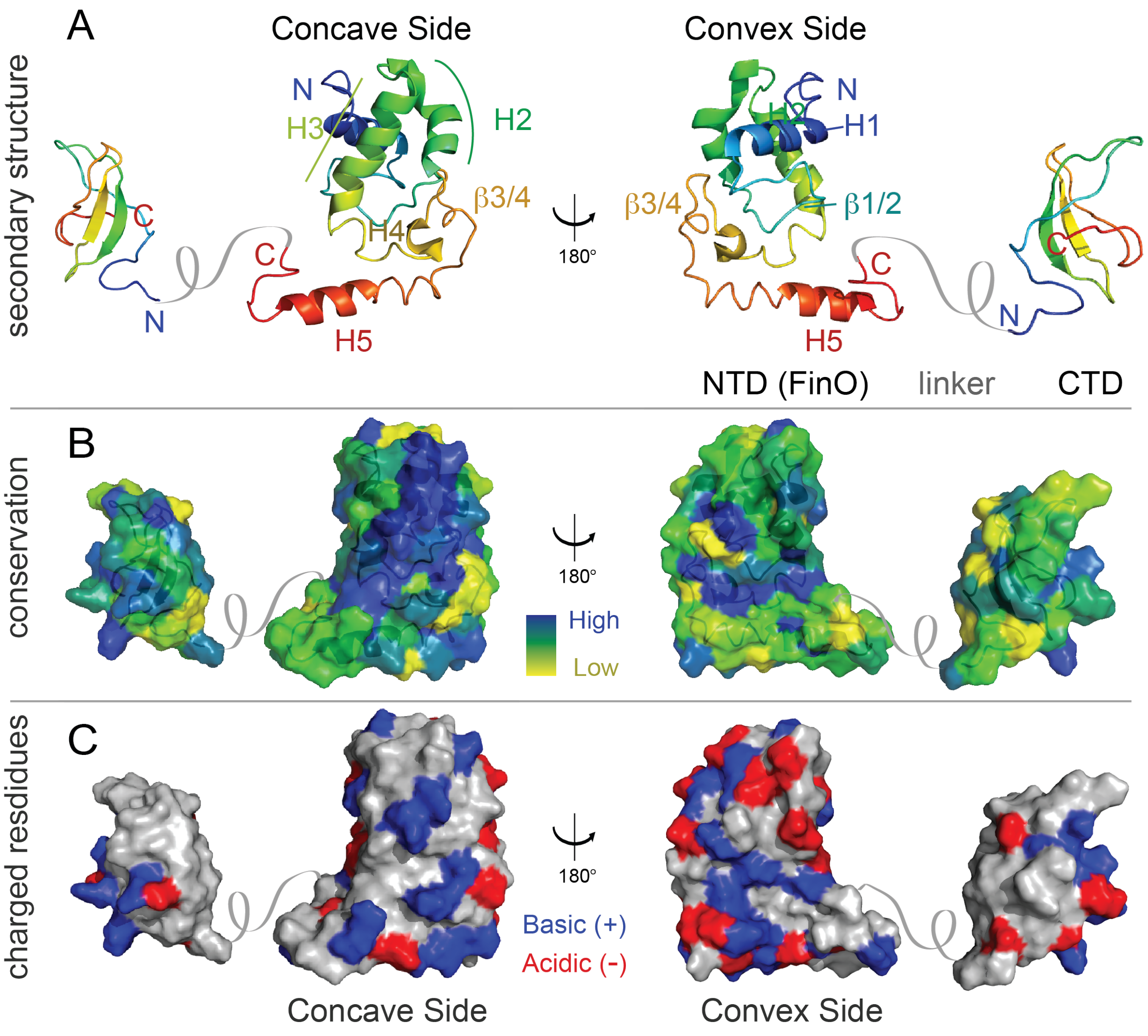


**Supplementary Figure 1. Structure of *E. coli* ProQ NTD and CTD.** (A) Cartoon representation of ProQ NTD (resi: 1-119, PDB ID: 5nb9) and CTD (resi: 180-232, PDB ID 5nbb) NMR structures (colored as rainbow chains (N-terminus: blue → C-terminus: red).(9) The less conserved linker domain (resi 120-179) is depicted as a hand-drawn grey coil. The NTD and CTD are shown in an arbitrary arrangement to one another. The right-hand side shows the same structures viewed from a 180° angle. N and C termini are indicated, as are the positions of key secondary-structure elements, as previously defined: α helices (H)1-5,(10) and β hairpins β1/2 and β3/4.(3) (B) Surface representations of ProQ NTD and CTD, colored according to degree of conservation (highly conserved: blue → minimally conserved: yellow). Conservation was determined by ConSurf server (8) using PDB ID 5nb9 as an input. (C) Surface representations of ProQ NTD and CTD, colored by electrostatic nature of side chains. Acidic residues (Asp and Glu), predicted to be negatively charged at cellular pH, are colored red; basic residues (Lys and Arg), predicted to be positively charged at cellular pH, are colored blue. Remaining surface residues are colored grey.

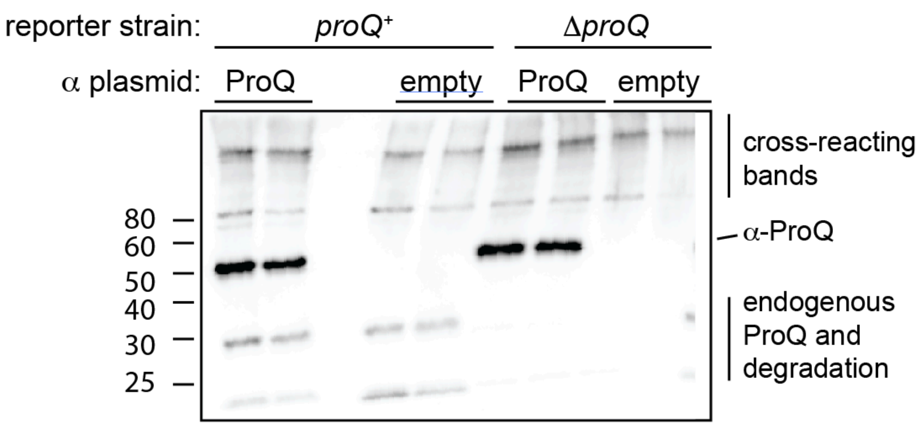


**Supplementary Figure 2. ProQ Western Blot and Identification of bands from B3H lysates.** Lysates were taken from β-galactosidase assays performed with *proQ*$^+$ (KB480) or Δ*proQ* (SP2) reporter strain cells containing three compatible plasmids: one the CI-MS2$^{CP}$ fusion protein, another that encoded α (empty) or the α-ProQ$^{ΔCTD}$ fusion protein (ProQ), and a third that encoded an MS2$^{hp}$-*cspE* hybrid RNA. Cells were grown in the presence of 0.2% arabinose and 10 μM IPTG, lysed, electrophoresed and blotted with an anti-ProQ polyclonal antibody (see Methods). The positions of molecular weight markers are indicated on the left. Assignments of bands to particular ProQ species are indicated on the right. Endogenous ProQ is expected to run at 26 kDa and the α-ProQ$^{ΔCTD}$ fusion protein at 48 kDa. The identities of these bands are further confirmed by their disappearance from lysates from a Δ*proQ* reporter (missing endogenous ProQ) or with an α-empty plasmid (missing α-ProQ$^{ΔCTD}$). The fact that the signal from α-ProQ is much higher than that arising from either cross-reacting bands or endogenous ProQ allowed for a dot-blot assay to meaningfully report on α-ProQ levels without prior electrophoresis (see Fig S7).

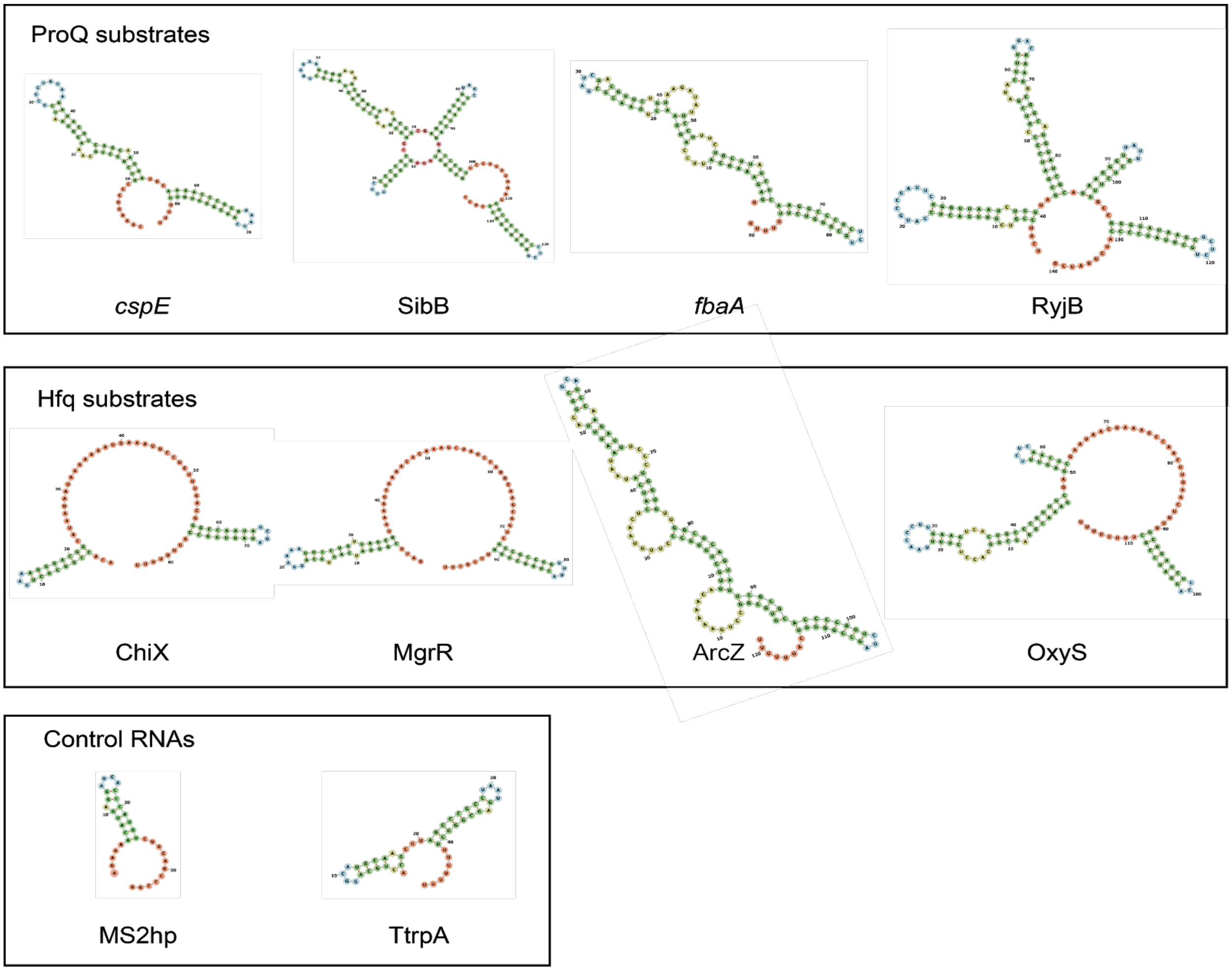


**Supplementary Figure 3. Predicted secondary structures of RNAs tested in B3H assay**. The predicted secondary structures of RNA sequences cloned into 1XMS2$^{hp}$-hybrid-RNA constructs and tested for B3H interactions in Fig 2. RNAs are separated according to whether they were selected for study based on literature evidence that they interact with ProQ (ProQ substrates), are Hfq-dependent sRNAs (Hfq substrates) or controls RNAs (see Results). Secondary structures were predicted with an RNA Fold algorithm and displayed using Forna.(6, 7)

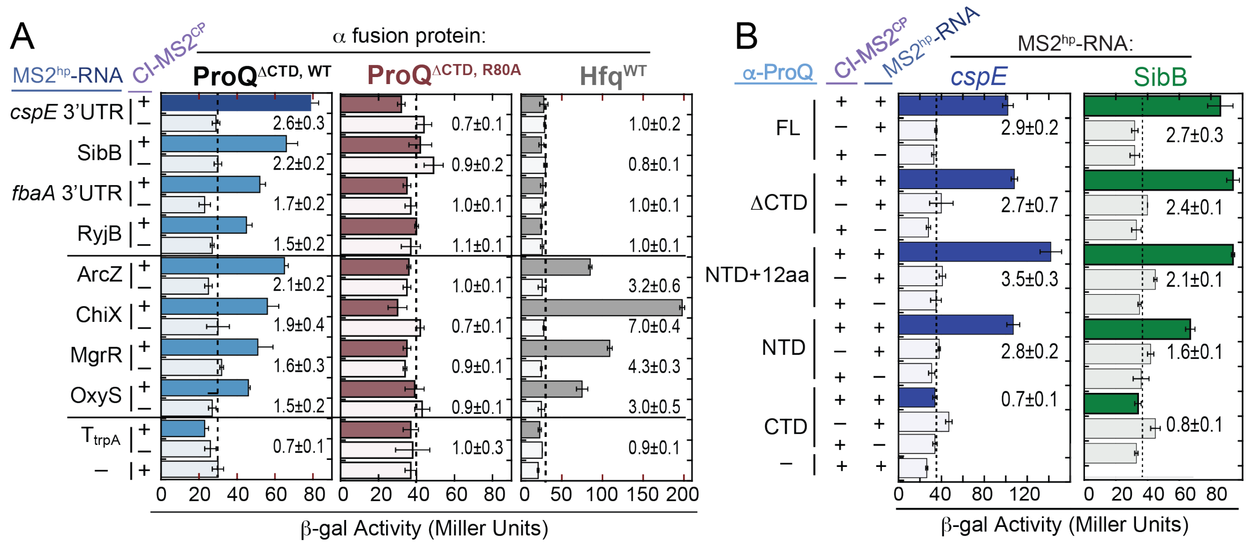


**Supplementary Figure 4. Absolute β-galactosidase values from representative datasets.** β-galactosidase (β-gal) values from which fold-stimulation over basal levels were calculated for data presented (A) Figure 2 and (B) Figure 3. (A) Results of a representative B3H assay between a panel of RNA substrates with wild type α-ProQ$^{WT\Delta CTD}$, α-ProQ$^{R80A\Delta CTD}$ or α-Hfq. β-gal assays were performed with *Δhfq* reporter strain cells containing three compatible plasmids: one that encoded λCI or the λCI-MS2$^{CP}$ fusion protein, another that encoded α or an α-fusion protein (α-ProQ$^{\Delta CTD}$, either with wild type ProQ or an R80A mutant, or α-Hfq), and a third that encoded a hybrid RNA (a single MS2$^{hp}$ moiety fused to *cspE* 3' UTR, SibB, *fbaA* 3' UTR, RyjB, ArcZ, ChiX, MgrR, OxyS, trpA terminator ($T_{trpA}$) or an RNA that contained only the MS2$^{hp}$ moiety. (B) Results of a representative B3H assay detecting interactions between α-ProQ truncations and *cspE* and SibB RNAs. β-gal assays were performed with *Δhfq* reporter strain cells containing three compatible plasmids: one that encoded λCI alone or the CI-MS2$^{CP}$ fusion protein, another that encoded α or an α-fusion protein (α-ProQ$^{FL}$ (full-length; resi=2-232), α-ProQ$^{\Delta CTD}$ (resi=2-176), α-ProQ$^{NTD+12aa}$ (resi=2-131), α-ProQ$^{NTD}$ (resi=2-119), or α-ProQ$^{CTD}$ (resi=181-232)), and a third that encoded a hybrid RNA (MS2$^{hp}$-*cspE* or MS2$^{hp}$-SibB) or an RNA that contained only the MS2$^{hp}$ moiety. For both panels, cells were grown in the presence of 0.2% arabinose and 50 µM IPTG. Bar graphs show absolute β-gal values as the averages and standard deviations of biological triplicate measurements. Each interaction is labeled with the fold-stimulation over basal levels (see Methods).

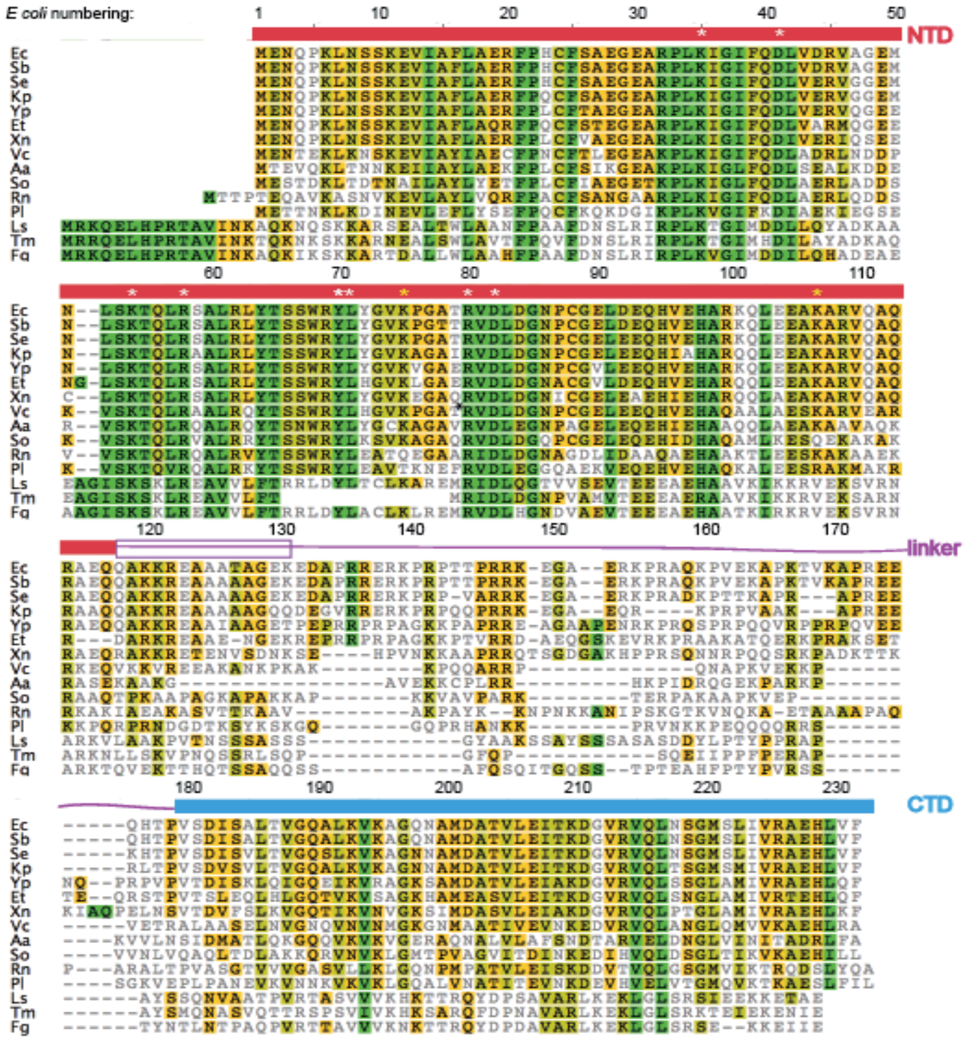


**Supplementary Figure 5. Multiple sequence alignment of ProQ and FinO sequences.** ProQ amino-acid sequences from the following species were obtained from NCBI: *Escherichia coli K12 (Ec), Shigella boydii (Sb), Salmonella enterica (Se), Klebsiella pneumoniae (Kp), Yersinia pestis (yp), Edwardsiella tardi (Et), Xenorhabdus nematophila (Xn), Vibrio cholerae (Vc), Aggregatibacter aphrophilus (Aa), Shewanella oneidensis (So), Rheinheimera nanhaiensis (Rn), Pseudoalteromonas luteoviolacea (Pl), Legionella shakespearei (Ls), Totlockia micdadei (Tm), Fluoribacter gormanii (Fg) Rheinheimera nanhaiensis*. Sequences were aligned using CLUSTALW with a BLOSUM cost matrix (reference). Residues are highlighted according to similarity (PAM120 score matrix; green, 100% similar; lime, 80-100% similar; yellow, 60% similar; no shading: 0-60% similar). Domain boundaries and residue numbers from *E. coli* ProQ are indicated above the sequences. Surface-exposed residues targeted for mutagenesis in this paper are indicated with an asterisk (white = highly conserved; yellow = less conserved). A purple box outlines the first 12 aa of the linker, which are included in the α-ProQ-NTD+12aa (resi=2-131) construct in Fig 3.

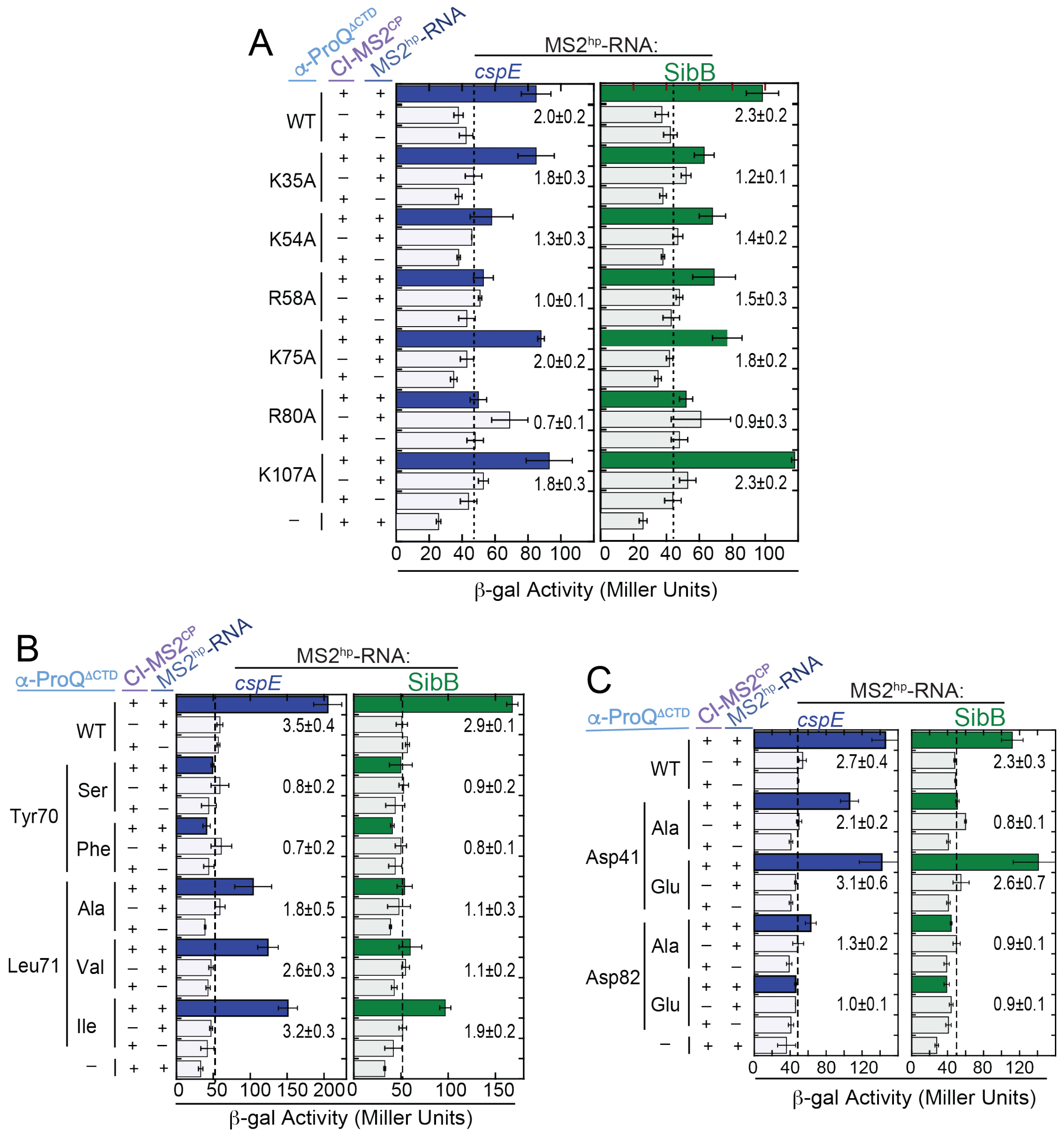


**Supplementary Figure 6. Absolute β-galactosidase values of representative datasets.** β-galactosidase (β-gal) values from which fold-stimulation over basal levels were calculated for data presented Figure 4. Results of a representative B3H assays showing effects on ProQ-RNA interactions of alanine mutations at (A) basic, (B) hydrophobic and (C) acidic residues. β-gal assays were performed with *Δhfq* reporter strain cells containing three compatible plasmids: one that encoded λCI or the λCI-MS2$^{CP}$ fusion protein, another that encoded the α-ProQ$^{ΔCTD}$ fusion protein (wild type (WT) or the indicated mutant), and a third that encoded a hybrid RNA (1XMS2$^{hp}$-*cspE* or 1XMS2$^{hp}$-SibB) or an RNA that contained only the MS2$^{hp}$ moiety. Cells were grown in the presence of 0.2% arabinose and 50 μM IPTG. Bar graphs show absolute β-gal values as the averages and standard deviations of biological triplicate measurements. Each interaction is labeled with the fold-stimulation over basal levels (see Methods).

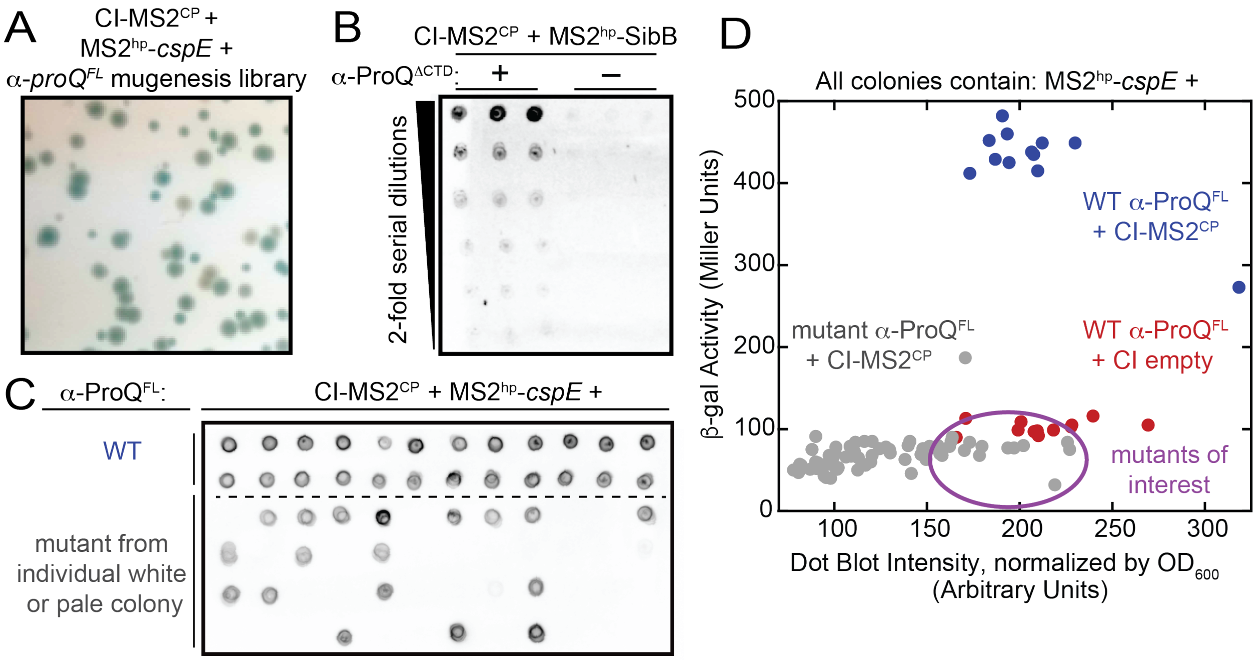


**Supplementary Figure 7. Primary and Counter Screen Process.** (A) Example of primary screen. Δ*hfq* reporter strain cells (SP5) were plated on LB agar medium containing 0.2% arabinose, 1.5 µM IPTG, 40 µg/mL X-gal and 125 µM TPEG. Cells contained three compatible plasmids: one that encoded the CI-MS2$^{CP}$ fusion protein, another that encoded an MS2$^{hp}$-*cspE* hybrid RNA, and a third that encoded a α-ProQ$^{FL}$ mutagenesis library (see Methods). Plates were grown overnight at 37°C and then incubated at 4°C. (B) Establishment of sensitivity and background of anti-ProQ dot-blot assay. Lysates were taken from β-galactosidase (β-gal) assays performed with KB511 reporter-strain cells containing three compatible plasmids: one the λCI-MS2$^{CP}$ fusion protein, another that encoded α (-) or the α-ProQ$^{ΔCTD}$ fusion protein (+), and a third that encoded an MS2$^{hp}$-SibB hybrid RNA. Cells were grown in the presence of 0.2% arabinose and 25 µM IPTG; two-fold dilutions of lysate were spotted on a nitrocellulose membrane and blotted with an anti-ProQ polyclonal antibody (see Methods (C) Representative dot blot from counterscreen. White and pale colonies from primary screen (individual white or pale colonies) or control colonies containing plasmids encoding wild-type α-ProQ$^{FL}$ along with CI-MS2$^{CP}$ and MS2$^{hp}$-*cspE* (WT) were picked and grown in the presence of 0.2% arabinose and 0 µM IPTG (see Methods). Cells were lysed for β-gal analysis and lysate from each well was spotted on nitrocellulose and blotted with anti-ProQ antibody, as in (A). Densitometry measurements were conducted in ImageJ. (D) For each well, arising from a single colony, β-gal activity was plotted versus densitometry-determined dot-blot intensity, normalized by the optical density ($OD_{600}$) of the well when lysed (see Methods). Plasmids were miniprepped and sequenced from colonies, indicated with a purple oval, that contained a α-*proQ*$^{FL}$ mutant sequence that gave rise to β-gal activity comparable to negative controls but anti-ProQ dot-blot intensity comparable to positive controls.

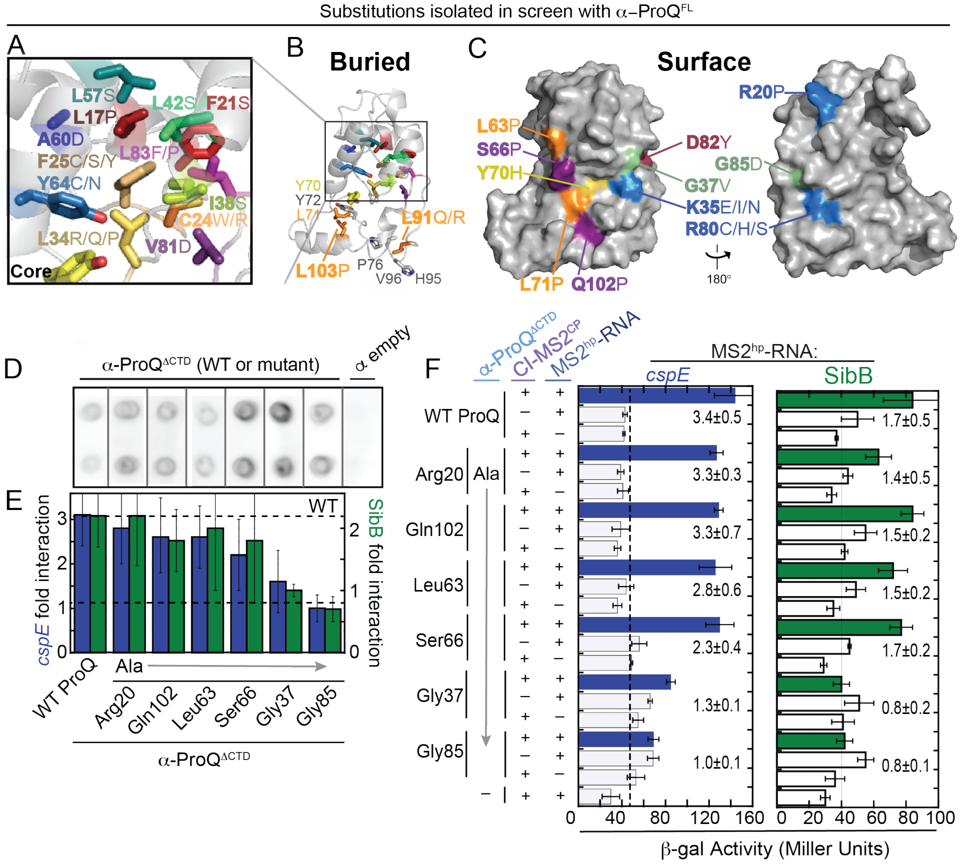


**Supplementary Figure 8. Results and validation of unbiased genetic screen.** (A) Surface and (B) cartoon representation of ProQ NTD structure (PDB ID: 5nb9),(9) viewed from concave face, showing buried residues at which substitutions were found to disrupt RNA binding in B3H screen with mutagenized α-ProQ$^{FL}$ plasmids (Table 1). Inset shows core residues identified in the RNA-binding screen. Each residue is shown in stick representation, and the substitution(s) identified in the screen is/are indicated with labels of matching color. (B) Additional buried residues at interface of H5 helix and core are shown as orange sticks and labeled with the substitution identified in the screen. Neighboring hydrophobic side chains not identified in the screen are shown as grey sticks. (C) Surface representation of ProQ NTD structure, viewed from concave face (left) and convex face (right), with positions of surface-exposed residues at which substitutions were found to disrupt RNA binding in B3H screen with mutagenized α-ProQ$^{FL}$ plasmids. Residue coloring: basic, blue; hydrophobic, orange; aromatic, yellow; acidic, red; polar: purple; glycine: green. Results of (D) dot-blot assay and (E) β-galactosidase (β-gal) assays testing the effects of alanine substitutions at residues implicated in screen. β-gal assays were performed with *Δhfq* reporter strain cells containing three compatible plasmids: one that encoded λCI or the CI-MS2$^{CP}$ fusion protein, another that encoded the α-ProQ$^{ΔCTD}$ fusion protein (wild-type, WT, or the indicated mutant), and a third that encoded a hybrid RNA (MS2$^{hp}$-*cspE* or MS2$^{hp}$-SibB) or an RNA that contained only the MS2$^{hp}$ moiety. The cells were grown in the presence of 0.2% arabinose and 50 μM IPTG. Lysates were spotted on nitrocellulose and blotted with anti-ProQ antibody, and also interrogated for β-gal activity (see Methods). The bar graph shows the fold-stimulation over basal levels as averages and standard deviations of values collected from three independent experiments conducted in triplicate across multiple days. (F) Absolute β-gal values of a representative dataset from which fold-stimulation over basal levels were calculated for data presented (E). Bar graphs show absolute β-gal values as the averages and standard deviations of biological triplicate measurements. Each interaction is labeled with the fold-stimulation over basal levels (see Methods).

A

| ***Ec* ProQ** | **G37** | **K35** | **D41** | **K54** | **R58** | **Y70** | **L71** | **R80** | **D82** | **G85** |
|---|---|---|---|---|---|---|---|---|---|---|
| **F' FinO** | G99 | ala | ala (-1: D102) | his (+1: K118) | R121 | Y133 | L134 | R143 | D145 | G148 |
| ***Nm* NMB1681** | G47 | ala | D51 | ala | ala (+1: R66) | Y77 | L78 | R87 | D89 | asn |
| ***Lp* RocC** | G52 | K50 | D56 | K71 | R75 | Y87 | L88 | R97 | D99 | G102 |
| ***Lp*1663** | G40 | K37 | D44 | K59 | R63 | Y75 | L76 | R85 | D87 | G90 |

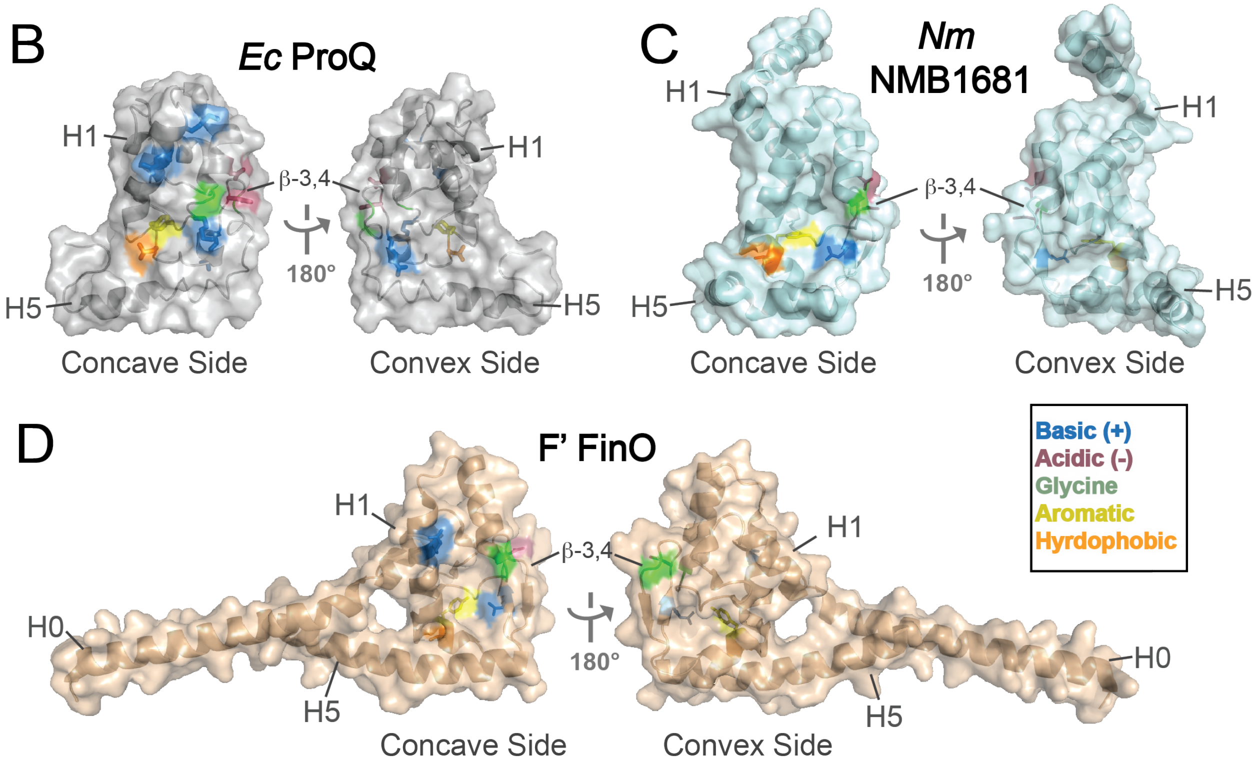


**Supplementary Figure 9. Conservation of *E. coli* ProQ RNA-binding residues across other FinO-domain proteins.** (A) The 10 residues identified in this work as important for RNA binding in *E. coli (Ec)* ProQ, whether from site-directed mutagenesis or a forward genetic screen, are listed in the top row, shaded by amino-acid character (basic, blue; hydrophobic, orange; aromatic, yellow; acidic, red; glycine: green). The corresponding residues in the *Ec* F' FinO protein, *N. meningitidis (Nm)* NMB1682, *L. pneumophila (Lp)* RocC and Lp1663 are listed in the rows below, based on a previous alignment. (10) If a residue is not conserved in the alignment, the amino acid in its place is listed. If a corresponding residue is present in the alignment at a position shifted by a single amino acid (-1/+1), that residue number is also indicated. Surface and cartoon representation of (B) *Ec* ProQ NTD NMR structure (PDB ID: 5nb9),(9) and crystal structures of (C) *Nm* NMB1682 (PDB ID: 3MW6),(5) or (D) *Ec* F' FinO protein (PDB ID: 1DVO),(3) viewed from concave face (left) and convex face (right), showing residues identified in this study as necessary for strong RNA interactions *in vivo,* or corresponding conserved residues, as listed in (A). Residues are shown in stick representation underneath a transparent surface, colored as above. While the conserved arginine residue at the position of Arg80 in both FinO and NM1681 has been modeled in crystal structures to be on the concave surface of the FinO-domain, in contrast to its position on the convex surface in the *Ec* ProQ NMR structure, this region is mediating crystal contacts in FinO and NM1681.(3, 5)

**Supplementary Table S1.** Plasmids used in this study. Listed in alphabetical order by plasmid name.

| Name | Description | Details | Reference/ Source | Figures |
|---|---|---|---|---|
| pACλCI | empty vector | Encodes full-length λCI under the control of the *lauUV5* promoter; confers CamR | (1) | throughout |
| pBrα | empty vector | Encodes residues 1-248 of alpha fused under the control of tandem *lpp* and *lacUV5* promoters; confers AmpR | (1) | throughout |
| pCH1 | pCDF-1XMS2$^{hp}$ | pCDF-pBAD-MS2hp-XmaI-HindIII; remove a 1xMS2 RNA hairpin from pKB845; confers SpcR | This study (oCH1, oCH2; see Methods) | throughout |
| pCH6 | pCDF-1XMS2$^{hp}$-ChiX | *E. coli chiX* cloned behind MS2hp in pCH1 between XmaI/HindIII sites; sRNA encodes its own terminator; confers SpcR | This study (oKB1190, oKB1991) | Fig 2 |
| pCH9 | pCDF-1XMS2$^{hp}$-OxyS | *E. coli oxyS* cloned behind MS2hp in pCH1 between XmaI/HindIII sites; sRNA encodes its own terminator; confers SpcR | This study (oKB1996, oKB1197) | Fig 2 |
| pCH10 | pCDF-1XMS2$^{hp}$-RhyB | *E. coli rhyB* cloned behind MS2hp in pCH1 between XmaI/HindIII sites; sRNA encodes its own terminator; confers SpcR | This study (oKB1098, oKB1099) | Fig 2 |
| pCH13 | pCDF-1XMS2$^{hp}$-ArcZ | *E. coli arcZ* cloned behind MS2hp in pCH1 between XmaI/HindIII sites; sRNA encodes its own terminator; confers SpcR | This study (oKB1211, oKB1212) | Fig 2 |
| pCW17 | pAC-$p_{constitutive}$-λCIMS2$^{CP}$ | Encodes residues1-248 of CI fused via an AAAEFPGIHPGM linker to a multimerization-resistant MS2 coat protein; transcription of fusion protein driven by a constitutive promoter; confers CamR | This study (see Methods) | throughout |
| pFW11-tet OL2 | pFW11tet-$O_L$2-62-lacZ | pFW11-derivative plasmid (2) bearing test promoter $p_{lac}O_L$2-62 fused to lacZ; confers TetR | Analogous to (4, 11) | N/A |
| pHL6 | pCDF-1XMS2$^{hp}$-$T_{trpA}$ | Intrinsic terminator from *trpA* operon cloned behind MS2hp in pCH1 between XmaI/HindIII sites; confers SpcR | This study (oHL19, oHL20; from pCH1) | Fig 2 |
| pKB817 | pBrα-Hfq | Encodes residues 1-248 of alpha fused via three alanine residues to full-length wild-type *E. coli* Hfq; confers AmpR | (11) | Fig 2C |
| pKB949 | pBrα-ProQ$^{FL}$ | Encodes residues 1-248 of alpha fused via three alanine residues to | This study (oKB1225, oKB1226) | Fig 3, Table 1 |

| | | | | |
|---|---|---|---|---|
| | | full-length wild-type *E. coli proQ*; confers AmpR | | |
| pKB951 | pBrα-ProQ$^{NTD+12aa}$ | Encodes residues 1-248 of alpha fused via three alanine residues to residues 1-131 of wild-type *E. coli proQ*; TAA stop codon after *proQ* added through PCR; confers AmpR | This study (oKB1225, oKB1227) | Fig 3 |
| pKB955 | pBrα-ProQ$^{\Delta CTD}$ | Encodes residues 1-248 of alpha fused via three alanine residues to residues 1-176 of wild-type *E. coli proQ*; TAA stop codon after *proQ* added through PCR; confers AmpR | This study (oKB1225, oKB1229) | throughout |
| pKB989 | pAC-plac-$\lambda$CIMS2$^{CP}$ | Encodes residues1-248 of CI fused via an AAAEFPGIHPGM linker to a multimerization-resistant MS2 coat protein; transcription of fusion protein driven by a lacUV5 promoter; confers CamR | (11) | N/A |
| pKB1067 | pFW11tet-$O_L$2-83-lacZ | pFW11-derivative plasmid(2) bearing test promoter $p_{lac}$ *$O_L$2-83-lacZ* ; confers TetR pFW11$_{tet}$, | This study (see Methods; oKB1362, oKB1363, oKB1366, oKB1367) | N/A |
| pSP10 | pCDF-1XMS2$^{hp}$-*cspE* 3'UTR | *E. coli* 3'UTR of *cspE* cloned behind MS2hp in pCH1 between XmaI/HindIII sites; mRNA encodes its own terminator; confers SpcR | This study (oSP10, oSP11) | throughout |
| pSP14 | pCDF-1XMS2$^{hp}$-SibB | *E. coli sibB* cloned behind MS2$^{hp}$ in pCH1 between XmaI/HindIII sites; sRNA encodes its own terminator; confers SpcR | This study (oSP16, oSP17) | throughout |
| pSP25 | pCDF-1XMS2$^{hp}$-RyjB | *E. coli ryjB* (140 nts) cloned behind MS2hp in pCH1 between XmaI/HindIII sites; confers SpcR | This study (oSP21, oSP23) | Fig 2 |
| pSP56 | pCDF-1XMS2$^{hp}$-*fbaA* 3'UTR | *E. coli* 3'UTR (terminal 91 nt of transcript, beginning +1038 after ATG) of *fbaA* cloned behind MS2$^{hp}$ in pCH1 between XmaI/HindIII sites; mRNA encodes its own terminator; confers SpcR | This study (oSP72, oSP76) | Fig 2 |
| pSP87 | pCDF-1XMS2$^{hp}$-MgrR | *E. coli mgrR* cloned behind MS2$^{hp}$ between XmaI/HindIII sites; sRNA its own terminator; confers SpcR | This study (oKB1109, oKB1110) | Fig 2 |
| pSP90 | pBrα-ProQ$^{NTD}$ | Encodes residues 1-248 of alpha fused via three alanine residues to residues 1-119 of wild-type *E. coli proQ*; TAA stop codon after *proQ* added through PCR; confers AmpR | This study (oKB1225, oSP126) | Fig 3 |

| | | | | |
|---|---|---|---|---|
| pSP92 | pBrα-ProQ$^{CTD}$ | Encodes residues 1-131 of alpha fused via three alanine residues to residues 181-232 of wild-type *E. coli proQ*; confers AmpR | This study (oSP128, oKB1226) | Fig 3 |
| **Plasmids cloned for site-directed mutagenesis** | | | | |
| pCG1 | pBrα-ProQ$^{\Delta CTD}$-K35A | *proQ* K35A mutation introduced to pKB955; confers AmpR | This study | Fig 4D |
| pCG3 | pBrα-ProQ$^{\Delta CTD}$-L71I | *proQ* L71I mutation introduced to pKB955; confers AmpR | This study (oSP90, oCG6) | Fig 4E |
| pSP111 | pBrα-ProQ$^{\Delta CTD}$-Y70F | *proQ* Y70F mutation introduced to pKB955; confers AmpR | This study (oSP87, oSP88) | Fig 4E |
| pSP112 | pBrα-ProQ$^{\Delta CTD}$-Y70S | *proQ* Y70S mutation introduced to pKB955; confers AmpR | This study (oSP87, oSP88) | Fig 4E |
| pSP114 | pBrα-ProQ$^{\Delta CTD}$-L71A | *proQ* L71A mutation introduced to pKB955; confers AmpR | This study (oSP90, oSP110) | Fig 4E |
| pSP113 | pBrα-ProQ$^{\Delta CTD}$-L71V | *proQ* L71V mutation introduced to pKB955; confers AmpR | This study (oSP89, oSP90) | Fig 4E |
| pSP101 | pBrα-ProQ$^{\Delta CTD}$-D41A | *proQ* D41A mutation introduced to pKB955; confers AmpR | This study (oSP77, oSP78) | Fig 4F |
| pSP102 | pBrα-ProQ$^{\Delta CTD}$-D41E | *proQ* D41E mutation introduced to pKB955; confers AmpR | This study (oSP77, oSP78) | Fig 4F |
| pSP104 | pBrα-ProQ$^{\Delta CTD}$-D82E | *proQ* D82E mutation introduced to pKB955; confers AmpR | This study (oSP79, oSP80) | Fig 4F |
| pSP103 | pBrα-ProQ$^{\Delta CTD}$-D82A | *proQ* D82A mutation introduced to pKB955; confers AmpR | This study (oSP79, oSP80) | Fig 4F |
| pSP107 | pBrα-ProQ$^{\Delta CTD}$-K54A | *proQ* K54A mutation introduced to pKB955; confers AmpR | This study (oSP83, oSP84) | Fig 4D |
| pSP137 | pBrα-ProQ$^{\Delta CTD}$-R58A | *proQ* R58A mutation introduced to pKB955; confers AmpR | This study (oSP86, oSP137) | Fig 4D |
| pSP136 | pBrα-ProQ$^{\Delta CTD}$-K75A | *proQ* K75A mutation introduced to pKB955; confers AmpR | This study (oSP134, oSP135) | Fig 4D |
| pSP117 | pBrα-ProQ$^{\Delta CTD}$-R80A | *proQ* R80A mutation introduced to pKB955; confers AmpR | This study (oSP92, oSP111) | Fig 1E,F Fig 2B, Fig 4D |
| pSP125 | pBrα-ProQ$^{\Delta CTD}$-G85A | *proQ* G85A mutation introduced to pKB955; confers AmpR | This study (oSP99, oSP100) | Fig S8D-F |
| pSP128 | pBrα-ProQ$^{\Delta CTD}$-K107A | *proQ* K107A mutation introduced to pKB955; confers AmpR | This study (oSP103, oSP104) | Fig 4D |
| pSP139 | pBrα-ProQ$^{\Delta CTD}$-Q102A | *proQ* Q102A mutation introduced to pKB955; confers AmpR | This study (oSP141, oSP142) | Fig S8D-F |
| pSP147 | pBrα-ProQ$^{\Delta CTD}$-R20A | *proQ* R20A mutation introduced to pKB955; confers AmpR | This study (oSP159, oSP160) | Fig S8D-F |
| pSP148 | pBrα-ProQ$^{\Delta CTD}$-S66A | *proQ* S66A mutation introduced to pKB955; confers AmpR | This study (oCG9, oCG10) | Fig S8D-F |
| pSP149 | pBrα-ProQ$^{\Delta CTD}$-G37A | *proQ* G37A mutation introduced to pKB955; confers AmpR | This study (oSP163, oSP132) | Fig S8D-F |
| pSP153 | pBrα-ProQ$^{\Delta CTD}$-L63A | *proQ* L63A mutation introduced to pKB955; confers AmpR | This study (oSP164, oSP165) | Fig S8D-F |

**Supplementary Table S2.** Strains used in this study.

| Name | Relevant Details | Antibiotic Resistance | Reference/ Source | Figures |
|---|---|---|---|---|
| NEB 5alpha F'Iq | *E. coli lacIq* host strain for plasmid construction | TetR | New England Biolabs | N/A |
| KB480 | FW102 harboring an F' episome bearing tetracycline resistance and test promoter *placO$_L$2-62* fused to *lacZ* | TetR, StrR | This study (see Methods) | Fig 1B, Fig S2 |
| KB483 | FW102 *hfq::kan* harboring F' episome bearing tetracycline resistance and test promoter *placO$_L$2-62* fused to *lacZ* | TetR, KanR, StrR | This study (see Methods) | throughout |
| KB496 | FW102 *hfq::kan* | KanR; StrR | (11) | N/A |
| KB511 | FW102 harboring an F' episome bearing tetracycline resistance and test promoter *placO$_L$2-82* fused to *lacZ*; | TetR, StrR | This study (see Methods) | N/A |
| SP2 | FW102 *proQ::kan* harboring an F' episome bearing tetracycline resistance and test promoter *placO$_L$2-62* fused to *lacZ* | TetR, KanR, StrR | This study (see Methods) | Fig 1C, Fig S2 |
| SP5 | FW102 *hfq::kan* harboring an F' episome bearing tetracycline resistance and test promoter *placO$_L$2-83* fused to *lacZ*; | TetR, KanR, StrR | This study (see Methods) | Genetic Screen (Table 1); results in Table S5 show verification in KB483. |

**Supplementary Table S3.** Oligonucleotides used in this study. M = C or A; Y = U or C; N = A/U/G/C

| Name | Description | Used for | Sequence |
|---|---|---|---|
| oCG1 | F ProQ D82A | Mutagenesis PCR: pSP103 | ACGCGTGTCGCTCTTGACGGC |
| oCG2 | R ProQ D82A | Mutagenesis PCR: pSP103 | TGCGCCGGGTTTAACACC |
| oCG3 | F ProQ K35A | Mutagenesis PCR: pCG1 | GCGTCCGCTGGCAATCGGTATTTTTC |
| oCG4 | R ProQ K35A | Mutagenesis PCR: pCG1 | GCTTCACCTTCCGCACTG |
| oCG6 | F ProQ L71I | Mutagenesis PCR: pCG3 | CTGGCGTTATATTTACGGTGTTA |
| oCG7 | F ProQ L91A | Mutagenesis PCR: pCG4 | ATGCGGTGAGGCGGACGAGCAAC |
| oCG9 | R ProQ L91A | Mutagenesis PCR: pCG4 | GGGTTGCCGTCAAGATCG |
| oCG9 | F ProQ S66A | Mutagenesis PCR: pSP148 | TCTCTACACTGCGAGCTGGCG |

| | | | |
|---|---|---|---|
| oCG10 | R ProQ S66A | Mutagenesis PCR: pSP148 | CGTAAAGCGGATCGCAATTG |
| oCH1 | F EcoRI | PCR: pCH1 | AATTCAGAAAACATGAGGATCACCCATGTCTGCAGC |
| oCH2 | R XmaI | PCR: pCH1 | CCGGGCTGCAGACATGGGTGATCCTCATGTTTTCTG |
| oCW6 | F pKB989 backbone | Gibson assembly: pCW17 | GCTAGCGCATGCCACACAGGAAACAGCGTATGAGCAC |
| oCW7 | R pKB989 backbone | Gibson assembly: pCW17 | CTCGAGGCCTGGGGTGCCTAATGAGTGAG |
| oCW18 | F constitutive promoter | Gibson assembly: pCW17 -10 | ACCCCAGGCCTCGAGTTTACGGCTAGCTCAGTCCTAGGTATAGTGCTAGCGCATGCCACA |
| oCW19 | R constitutive promoter | Gibson assembly: pCW17 -10 | TGTGGCATGCGCTAGCACTATACCTAGGACTGAGCTAGCCGTAAACTCGAGGCCTGGGGT |
| oCW28 | F pCW17 -35 | Mutagenesis PCR: pCW17 | CCAGGCCTCGAGACGATAGCTAGCTCAGTC |
| oCW29 | R pCW17 -35 | Mutagenesis PCR: pCW17 | GACTGAGCTAGCTATCGTCTCGAGGCCTGG |
| oHL19 | F $T_{TrpA}$ | Mutagenesis PCR: pHL6 | AGCGGGCTTTTTTTTCAGCTTGGCTGTTTTGGCGGATGAG |
| oHL20 | R $T_{trpA}$ | Mutagenesis PCR: pHL6 | CATTAGGCGGGCTAAGCTTGCATGCCTGCAGGTCCCGGGC |
| oKB1190 | F XmaI ChiX | PCR: pCH6 | CGCCAATAGCGATATTGGCCATTTTTTTAAGCTTGGCGG |
| oKB1191 | R HindIII ChiX | PCR: pCH6 | CCGGCCAAGCTTAAAAAAATGGCCAATATCGCTATTGGC |
| oKB1198 | F XmaI RhyB | PCR: pCH10 | TCCCCCCGGGGCGATCAGGAAGACCCTCG |
| oKB1199 | R HindIII RhyB | PCR: pCH10 | CCGGCCAAGCTTAAAAAAAAGCCAGCACCCGG |
| oKB1196 | F XmaI OxyS | PCR: pCH9 | TCCCCCCGGGGAAACGGAGCGGCACCTC |
| oKB1197 | R HindIII OxyS | PCR: pCH9 | CCGGCCAAGCTTAAAAAAAAGCGGATCCTGGAGATCC |
| oKB1211 | F XmaI ArcZ | PCR: pCH13 | TCCCCCCGGGGTGCGGCCTGAAAAACAGTGC |
| oKB1212 | R HindIII ArcZ | PCR: pCH13 | CCGGCCAAGCTTAAAAAATGACCCCGGCTAGACC |
| oKB1192 | F XmaI McaS | PCR: pSP59 | TCCCCCCGGGACCGGCGCAGAGGAG |
| oKB1193 | R HindIII McaS | PCR: pSP59 | CCGGCCAAGCTTAAAAAATAGAGTCTGTCGACATCCGCC |
| oKB1109 | F XmaI MgrR | PCR: pSP87 | TCCCCCCGGGGATTCGTTATCAGTGCAGGAAAATGCC |
| oKB1110 | R HindIII MgrR | PCR: pSP87 | CCGGCCAAGCTTAAAAAAAACCGCCAGTAAACCGGC |

| | | | |
|---|---|---|---|
| oKB1225 | F NotI *proQ* | PCR: pKB949, pKB955, pSP90, pKB951 | ATAAGAATGCGGCCGCAGAAAATCAACCTAAGTTGAATAGCAGTAAAG |
| oKB1226 | R BamHI *proQ* | PCR: pKB949, pSP92 | CTTCGGATCCTCAGAACACCAGGTGTTCTGCGC |
| oKB1227 | R BamHI *proQ* NTD+12aa | PCR: pKB951 | CTTCGGATCCTTATTTCTCACCAGCAGTTGCG |
| oKB1229 | R BamHI *proQ* ∆CTD | PCR: pKB955 | CTTCGGATCCTTACTGTTCTTCGCGAGGTGCTTTTAC |
| oKB1362 | F EcoRI | PCR: pKB1067 | GGCCGGGAATTCTTCCACCGGCGG |
| oKB1363 | R HindIII | PCR: pKB1067 | CCGGCCAAGCTTGGCTGCAGGTCG |
| oKB1366 | F overlap +21bp | PCR: pKB1067 | CAACACCGCCAGAGATAGCTGCCACGGTGCCCGACCGTCCTGAGGCACCCCGGGC |
| oKB1367 | R overlap +21bp | PCR: pKB1067 | GCCCGGGGTGCCTCAGGACGGTCGGGCACCGTGGCAGCTATCTCTGGCGGTGTTG |
| oSP10 | F XmaI *cspE* 3'UTR | PCR: pSP93, pSP10 | GGCCGGCCCGGGCAAAGGCCCTTCTGCTGCAAAC |
| oSP11 | R HindIII *cspE* 3'UTR | PCR: pSP93, pSP10 | CCGGCCAAGCTTAAAAAAAACCCGCTGATTAAGCGGGT |
| oSP16 | F XmaI SibB | PCR: pSP14 | GGCCGGCCCGGGGAGGGTAGAGCGGGGTTTCC |
| oSP17 | R HindIII SibB | PCR: pSP14 | CCGGCCAAGCTTGGAAAGCCCCTCCCGAGG |
| oSP21 | F XmaI RyjB | PCR: pSP25 | GGCCGGCCCGGGTCATCCGTCGTTGACTCCATGCC |
| oSP23 | R HindIII RyjB | PCR: pSP25 | CCGGCCAAGCTTCGATAAGATGGGGATAAGCAGAGCGCTTAT |
| oSP76 | F XmaI *fbaA* 3'UTR | PCR: pSP56 | GGCCGGCCCGGGGGAGAAAGCATTCCAGGAACTGAACG |
| oSP72 | R HindIII *fbaA* 3'UTR | PCR: pSP56 | CCGGCCAAGCTTAAAAAAAGACCCGCAGAGCGGG |
| oSP77 | F ProQ D41 (A/E) | Mutagenesis PCR: pSP101, pSP102 | ATTTTTCAGGMGTTGGTCGATCGTGTTGC |
| oSP78 | R ProQ D41 (A/E) | Mutagenesis PCR: pSP101, pSP102 | ACCGATTTTCAGCGGACG |
| oSP79 | F ProQ D82 (A/E) | Mutagenesis PCR: pSP104 | AACGCGTGTCGMGCTTGACGGCAAC |
| oSP80 | R ProQ D82 (A/E) | Mutagenesis PCR: pSP104 | GCGCCGGGTTTAACACCG |
| oSP83 | F ProQ K54 (A/E) | Mutagenesis PCR: pSP107 | GAACCTGAGCGMGACGCAATTGCG |
| oSP84 | R ProQ K54 (A/E) | Mutagenesis PCR: pSP107, pSP145 | ATTTCCCCAGCAACACGA |
| oSP86 | R ProQ R58A | Mutagenesis PCR: pSP137, pSP150 | TTGCTCAGGTTCATTTCC |
| oSP87 | F ProQ Y70 (F/S) | Mutagenesis PCR: pSP111, pSP112 | GAGCTGGCGTTYTCTTTACGGTG |

| | | | |
|---|---|---|---|
| oSP88 | R ProQ Y70 (F/S) | Mutagenesis PCR: pSP111, pSP112 | GAAGTGTAGAGACGTAAAGC |
| oSP89 | F ProQ L71 (A/V) | Mutagenesis PCR: pSP113, | CTGGCGTTATGNTTACGGTGTTAAAC |
| oSP90 | R ProQ L71 (A/V) | Mutagenesis PCR: pSP113, pSP114, pCG3 | CTCGAAGTGTAGAGACGTAAAG |
| oSP92 | R ProQ R80A | Mutagenesis PCR: pSP117 | GGTTTAACACCGTAAAGATAAC |
| oSP99 | F ProQ G85 (D/A) | Mutagenesis PCR: pSP125 | CGATCTTGACGMTAACCCATGCGGTGAGCTGGAC |
| oSP100 | R ProQ G85 (D/A) | Mutagenesis PCR: pSP125 | ACACGCGTTGCGCCGGGT |
| oSP103 | F ProQ K107 (A/E) | Mutagenesis PCR: pSP128 | TGAAGAAGCGGMGGCGCGTGTTCAGG |
| oSP104 | R ProQ K107 (A/E) | Mutagenesis PCR: pSP128 | AGCTGCTTGCGAGCATGC |
| oSP110 | F ProQ L71A | Mutagenesis PCR: pSP114 | CTGGCGTTATGCTTACGGTGTTAAAC |
| oSP111 | F ProQ R80A | Mutagenesis PCR: pSP117, pSP151, pSP152, pSP160 | CGGCGCAACGGCGGTCGATCTTG |
| oSP126 | R BamHI proQ NTD | PCR: pSP90 | CCGGCCGGATCCTTACGCTTGCTGTTCAGCACG |
| oSP128 | F NotI I *proQ* CTD | PCR: pSP92 | GAATGCGGCCGCATCTGACATTTCAGCTCTGACTGTCGGA |
| oSP135 | F ProQ K75A | Mutagenesis PCR: pSP136 | TTACGGTGTTGCACCCGGCGCAA |
| oSP134 | R ProQ K75A | Mutagenesis PCR: pSP136 | AGATAACGCCAGCTCGAAG |
| oSP137 | F ProQ R58A | Mutagenesis PCR: pSP137 | AACGCAATTGGCATCCGCTTTAC |
| oSP141 | F ProQ Q102A | Mutagenesis PCR: pSP137 | TGCTCGCAAGGCGCTTGAAGAAG |
| oSP142 | R ProQ Q102A | Mutagenesis PCR: pSP137 | TGCTCTACATGTTGCTCG |
| oSP163 | F ProQ G37A | Mutagenesis PCR: pSP149 | GCTGAAAATCGCTATTTTTCAGGATTTGGTCGATCG |
| oSP132 | R ProQ G37A | Mutagenesis PCR: pSP149 | GGACGCGCTTCACCTTCC |
| oSP159 | F ProQ R20A | Mutagenesis PCR: pSP147 | TCTGGCCGAAGCTTTTCCCCACTG |
| oSP160 | R ProQ R20A | Mutagenesis PCR: pSP147 | AACGCGATTACTTCTTTAC |
| oSP164 | F ProQ L63A | Mutagenesis PCR: pSP153 | CGCTTTACGTGCCTACACTTCGAG |
| oSP165 | R ProQ L63A | Mutagenesis PCR: pSP153 | GATCGCAATTGCGTTTTG |

**Supplementary Table S4. Validation of dot-blot assay as an effective counter-screen to identify pBRα-*proQ* plasmids encoding α-ProQ$^{FL}$ fusion proteins with low expression levels.** B3H reporter-strain cells (SP5) containing the CI-MS2$^{CP}$ adapter protein and either the MS2$^{hp}$-*cspE* hybrid RNA were transformed with an pBrα-*proQ$^{FL}$* mutagenesis library and plated appropriate on indicator medium to enable a clear distinction between blue positive-control colonies (containing the WT fusion proteins and the *cspE* hybrid RNA) and white negative-control colonies (instead containing a plasmid encoding α-empty). Ten white or pale colonies were picked, grown in liquid culture and ProQ-expression levels were determined by dot-blot assay and densitometry (see Methods and Fig S7). Plasmids were mini-prepped from 5 colonies with ProQ levels similar to those with an α-empty plasmid and 5 colonies with ProQ levels similar to WT α-ProQ controls. The *proQ* sequence from pBrα-*proQ$^{FL}$* plasmids in each colony was sequenced and compared to WT *proQ*. The consequences of each mutation present are indicated in terms of whether they lead to a premature stop codon and the location of each amino-acid substitution, including whether a mutation alters a codon to an in-frame stop codon (X) or introduces a frameshift (fs) that leads to stop codon (fsX) shortly thereafter.

| ProQ expression levels | Mutant Name | Premature Stop Codon? | Consequence of Mutation |
|---|---|---|---|
| Low Levels (~α-empty) | 0Mn-48 | Yes | L42fsX |
| | 0Mn-57 | Yes | L61X |
| | 0Mn-62 | Yes | Q40fsX |
| | 0Mn-67 | Yes | N3fsX |
| | 0Mn-89 | Yes | Q40fsX |
| High Levels (~WT α-ProQ$^{FL}$) | 0Mn-52 | No | Y70H |
| | 0Mn-53 | No | S66P |
| | 0Mn-68 | No | F25S |
| | 0Mn-100 | No | I38T; V43D |
| | 0Mn-101 | No | A15T; D82Y |

**Supplementary Table S5. Validation and Quantification of Results from Unbiased Screen**. Results of β-galactosidase (β-gal) and dot-blot assays to confirm behavior of pBrα-*proQ*$^{FL}$ plasmids isolated in forward genetic screen in KB483 reporter cells (carrying *$O_L$2-62-lacZ* reporter). Reporter-strain KB483 cells containing KB483 containing plasmids encoding *(i)* the CI-MS2$^{CP}$ fusion protein, and *(ii)* either a MS2$^{hp}$-*cspE (cspE)* or MS2$^{hp}$-SibB (SibB) hybrid RNA were transformed with plasmids isolated from white/pale primary-screen colonies that passed the dot-blot counter-screen (see Methods) in order to introduce the mutant pBRα-*proQ*$^{FL}$ plasmid contained in each colony. Cells were grown in triplicate in the presence of 0.2% arabinose and either 0 µM or 50 µM IPTG. Lysates from colonies containing MS2$^{hp}$-*cspE* were then assessed via dot-blot assay with an anti-ProQ antibody and densitometry was conducted to quantify the intensity of each dot. β-gal values from triplicate colonies were averaged and then normalized to the value from comparable colonies containing pBRα-empty as a negative control. This fold-stimulation value from each *proQ* mutant, along with average dot-blot intensity was then normalized to the values arising from WT α-ProQ (set to 1.0) and from α-empty (set to 0.0) in cells grown at 0 µM IPTG. The three values that were set to 1.0 are indicated in bold text within the table. Thus, the normalized expression levels (dot-blot intensity) and fold-stimulation of each mutant can be directly compared between IPTG concentrations. A small number of α–ProQ variants show lower expression levels than WT when both are induced with 0 µM IPTG, but induction of each variant with 50 µM IPTG leads to higher expression levels than WT α–ProQ at 0 µM IPTG. For each of these α–ProQ variants, induction with 50 µM IPTG, nevertheless results in lower B3H interaction for the variant than WT α–ProQ in the absence of IPTG. Errors reported for both sets of measurements represent one standard deviation from each mean value, propagated through each normalization step. For a description of variables listed under "Mutant Information," see Table 1.

| Mutant Information | | | | | Normalized β-gal Fold-Stimulation | | | | Normalized Dot-Blot Intensity | |
|---|---|---|---|---|---|---|---|---|---|---|
| Resi | Loc | pBRα-*proQ*$^{FL}$ Mutant Name | RNA | Variant | 0 µM IPTG | | 50 µM IPTG | | 0 µM IPTG | 50 µM IPTG |
| | | | | | *cspE* | SibB | *cspE* | SibB | | |
| | | | | WT | **1.0±0.2** | **1.0±0.2** | 0.9±0.2 | 0.9±0.4 | **1.0±0.4** | 2.7±1.2 |
| | | | | α empty | 0.0 | 0.0 | 0.0 | 0.0 | **0.0** | 0.0 |
| L17 | core | 0-Mn-56; 0.1Mn-SibB-55 | both | L17P | 0.2±0.1 | 0.2±0.2 | 0.1±0.1 | 0.3±0.1 | 0.7±0.4 | 1.4±0.5 |
| R20 | surface | 0.1Mn-SibB-49 | SibB | R20P | 0.2±0.1 | 0.2±0.2 | 0.0±0.1 | 0.2±0.1 | 0.8±0.5 | 1.0±0.5 |
| F21 | core | 0Mn-SibB-27; 0.1Mn-SibB-5 | SibB | F21S | 0.3±0.1 | 0.1±0.4 | 0.1±0.1 | 0.3±0.1 | 1.7±0.6 | 3.0±1.2 |
| C24 | core | 0-Mn-55 | *cspE* | C24W | 0.2±0.1 | 0.2±0.2 | 0.1±0.1 | 0.3±0.1 | 0.7±0.3 | 2.7±1.9 |
| | | 0.1Mn-SibB-16 | SibB | C24R | 0.2±0.1 | 0.2±0.2 | 0.1±0.1 | 0.5±0.2 | 1.0±0.4 | 3.4±1.2 |
| F25 | core | 0.1Mn-50 | *cspE* | F25C | 0.2±0.1 | 0.2±0.2 | 0.1±0.1 | 0.4±0.2 | 0.9±0.6 | 2.0±1.5 |
| | | 0-Mn-68; 0.1Mn-60; 0Mn-SibB-24 | both | F25S | 0.3±0.1 | 0.2±0.2 | 0.2±0.1 | 0.3±0.1 | 2.1±0.8 | 3.7±2.9 |
| | | 0.1Mn-SibB-82 | SibB | F25Y | 0.4±0.1 | 0.3±0.2 | 0.3±0.1 | 0.3±0.1 | 1.3±0.5 | 2.6±2.4 |
| L34 | core | 0-Mn-44 | *cspE* | L34R | 0.2±0.1 | 0.2±0.2 | 0.2±0.1 | 0.3±0.2 | 1.3±0.7 | 4.5±1.7 |
| | | 0.1Mn-SibB-34 | SibB | L34Q | 0.2±0.1 | 0.2±0.2 | 0.1±0.1 | 0.3±0.1 | 0.7±0.5 | 2.1±1.5 |
| | | 0.1Mn-SibB-50 | SibB | L34P | 0.3±0.1 | 0.2±0.2 | 0.1±0.1 | 0.2±0.1 | 1.7±0.9 | 3.6±2.2 |

| | | | | | | | | | | |
|---|---|---|---|---|---|---|---|---|---|---|
| K35 | surface | 0Mn-SibB-23 | SibB | K35E | 0.2±0.1 | 0.3±0.2 | 0.1±0.1 | 0.3±0.1 | 1.2±0.7 | 1.4±1.4 |
| | | 0Mn-SibB-5 | SibB | K35N | 0.3±0.1 | 0.2±0.2 | 0.2±0.1 | 0.3±0.1 | 0.8±0.3 | 3.0±1.4 |
| | | 0.1Mn-SibB-66 | SibB | K35I | 0.3±0.1 | 0.2±0.2 | 0.1±0.1 | 0.3±0.1 | 1.3±0.8 | 3.1±1.7 |
| G37 | surface | 0Mn-SibB-17 | SibB | G37V | 0.3±0.1 | 0.1±0.2 | 0.1±0.1 | 0.2±0.1 | 2.0±1.0 | 2.8±1.1 |
| I38 | core | 0-Mn-58 | *cspE* | I38S | 0.3±0.1 | 0.2±0.2 | 0.0±0.1 | 0.3±0.1 | 1.3±0.9 | 1.5±0.7 |
| L42 | core | 0Mn-SibB-75 | SibB | L42S | 0.3±0.1 | 0.2±0.2 | 0.1±0.1 | 0.3±0.1 | 1.1±0.6 | 3.5±1.3 |
| L57 | core | 0-Mn-14 | *cspE* | L57S | 0.3±0.1 | 0.2±0.2 | 0.1±0.1 | 0.2±0.1 | 1.1±0.4 | 3.5±1.7 |
| A60 | core | 0-Mn-122 | *cspE* | A60D | 0.2±0.1 | 0.1±0.2 | 0.1±0.1 | 0.2±0.1 | 1.3±0.5 | 2.4±1.3 |
| L63 | surface | 0.1Mn-SibB-60 | SibB | L63P | 0.3±0.1 | 0.3±0.2 | 0.1±0.1 | 0.1±0.1 | 1.0±0.5 | 1.5±0.6 |
| Y64 | core | 0-Mn-99 | *cspE* | Y64C | 0.2±0.1 | 0.2±0.2 | 0.1±0.1 | 0.2±0.1 | 1.5±0.5 | 2.7±0.9 |
| | | 0-Mn-136; 0Mn-SibB-61 | both | Y64N | 0.2±0.1 | 0.3±0.2 | 0.1±0.1 | 0.1±0.1 | 1.0±0.5 | 0.9±0.4 |
| S66 | surface | 0.1Mn-SibB-44; 0-Mn-53 | both | S66P | 0.2±0.1 | 0.2±0.2 | 0.1±0.1 | 0.1±0.1 | 1.0±0.3 | 1.9±0.6 |
| Y70 | surface | 0-Mn-52; 0.1Mn-61; 0Mn-SibB-8 | both | Y70H | 0.3±0.1 | 0.2±0.2 | 0.1±0.1 | 0.1±0.1 | 1.8±1.1 | 2.9±2.1 |
| L71 | surface | 0-Mn-71; 0.1Mn-80; 0Mn-SibB-40 | both | L71P | 0.3±0.1 | 0.2±0.2 | 0.1±0.1 | 0.1±0.1 | 1.8±1.0 | 1.5±0.6 |
| R80 | surface | 0-Mn-39 | *cspE* | R80C | 0.2±0.1 | 0.3±0.2 | 0.1±0.1 | 0.3±0.1 | 2.4±0.9 | 2.8±1.2 |
| | | 0-Mn-98 | *cspE* | R80H | 0.4±0.1 | 0.3±0.2 | 0.1±0.1 | 0.2±0.1 | 2.9±1.3 | 4.4±1.7 |
| | | 0.1Mn-SibB-43 | SibB | R80S | 0.2±0.1 | 0.2±0.2 | 0.1±0.1 | 0.2±0.1 | 1.0±0.4 | 1.7±0.8 |
| V81 | core | 0-Mn-64; 0.1Mn-97; 0.1Mn-SibB-23 | both | V81D | 0.2±0.1 | 0.3±0.2 | 0.1±0.1 | 0.3±0.1 | 1.4±0.8 | 2.3±1.0 |
| D82 | surface | 0.1Mn-151 | *cspE* | D82Y | 0.2±0.1 | 0.3±0.2 | 0.2±0.1 | 0.3±0.1 | 2.5±0.9 | 3.0±1.0 |
| L83 | core | 0-Mn-158 | *cspE* | L83F | 0.2±0.1 | 0.3±0.2 | 0.3±0.1 | 0.2±0.1 | 2.2±1.6 | 2.0±1.2 |
| | | 0Mn-SibB-57 | SibB | L83P | 0.3±0.1 | 0.2±0.2 | 0.3±0.1 | 0.2±0.1 | 1.3±0.8 | 1.9±1.0 |
| G85 | surface | 0.1Mn-164; 0.1Mn-SibB-20 | both | G85D | 0.2±0.1 | 0.2±0.2 | 0.3±0.1 | 0.2±0.1 | 2.2±0.9 | 2.5±1.2 |
| L91 | buried | 0Mn-SibB-32 | SibB | L91Q | 0.2±0.1 | 0.3±0.2 | 0.2±0.1 | 0.2±0.1 | 4.2±1.6 | 3.3±1.1 |
| | | 0-Mn-113 | *cspE* | L91R | 0.1±0.1 | 0.3±0.2 | 0.2±0.1 | 0.3±0.1 | 2.9±1.2 | 2.2±0.7 |
| Q102 | surface | 0Mn-SibB-78 | SibB | Q102P | 0.1±0.1 | 0.3±0.2 | 0.1±0.1 | 0.3±0.1 | 0.7±0.5 | 1.2±0.8 |
| L103 | buried | 0.1Mn-SibB-32 | SibB | L103P | 0.2±0.1 | 0.3±0.2 | 0.3±0.1 | 0.3±0.1 | 3.5±1.2 | 2.8±1.1 |
